# The structural role of SARS-CoV-2 genetic background in the emergence and success of spike mutations: the case of the spike A222V mutation

**DOI:** 10.1101/2021.12.05.471263

**Authors:** Tiziana Ginex, Clara Marco-Marín, Miłosz Wieczór, Carlos P. Mata, James Krieger, Maria Luisa López-Redondo, Clara Francés-Gómez, Paula Ruiz-Rodriguez, Roberto Melero, Carlos Óscar Sánchez-Sorzano, Marta Martínez, Nadine Gougeard, Alicia Forcada-Nadal, Sara Zamora-Caballero, Roberto Gozalbo-Rovira, Carla Sanz-Frasquet, Jeronimo Bravo, Vicente Rubio, Alberto Marina, The IBV-Covid19-Pipeline, Ron Geller, Iñaki Comas, Carmen Gil, Mireia Coscolla, Modesto Orozco, José Luis Llácer, José-Maria Carazo

**Affiliations:** Centro de Investigaciones Biológicas Margarita Salas (CIB-CSIC), 28040 Madrid, Spain; Instituto de Biomedicina de Valencia (IBV-CSIC), 46010 Valencia, Spain; Molecular Modeling and Bioinformatics, Institute for Research in Biomedicine (IRB Barcelona), The Barcelona Institute of Science and Technology (BIST), 08036 Barcelona, Spain; Centro Nacional de Biotecnología (CNB-CSIC), 28049 Madrid, Spain; I2SysBio, University of Valencia-CSIC, FISABIO Joint Research Unit Infection and Public Health, 46020 Valencia, Spain; Centro para Investigación Biomédica en Red sobre Epidemiología y Salud Pública (CIBERESP), 28029 Valencia, Spain; Centro de Investigación Biomédica en Red en Enfermedades Raras (CIBERER), 28029 Madrid, Spain; Centro Nacional de Microbiología (CNM-ISCIII), Instituto de Salud Carlos III, 28220 Madrid, Spain

**Author notes:** Correspondence to José-Maria Carazo. These authors contributed equally (Tiziana Ginex, Clara Marco-Marín, Miłosz Wieczór, Carlos P. Mata, James Krieger).

## Abstract

The S:A222V point mutation, within the G clade, was characteristic of the 20E (EU1) SARS-CoV-2 variant identified in Spain in early summer 2020. This mutation has now reappeared in the Delta subvariant AY.4.2, raising questions about its specific effect on viral infection. We report combined serological, functional, structural and computational studies characterizing the impact of this mutation. Our results reveal that S:A222V promotes an increased RBD opening and slightly increases ACE2 binding as compared to the parent S:D614G clade. Finally, S:A222V does not reduce sera neutralization capacity, suggesting it does not affect vaccine effectiveness.

## Main

Since the start of the COVID19 pandemic in early 2020 the causative agent, SARS-CoV-2, has been diversifying. Hundreds of variants have been identified, associated with thousands of mutations. Most of these mutations have no assigned role and they thrive in the population due to different stochastic population genetics forces such as founder or superspreading events (van Dorp et al. 2020). However, a handful of variants are known to have evolved increased transmissibility and/or ability to reduce antibody responses. Most of those variants are characterized by a constellation of mutations, from five to more than 15, in the spike (S) protein. While some mutations have been deeply characterized, the role of most of these mutations in underlying the phenotypes associated with the variants and their mechanisms of action remain unknown.

Furthermore, many of the mutations found in the most successful variants, including Alpha, Delta or Beta, have been seen before in other genetic backgrounds, usually in isolation, where they have not been able to thrive in the population.

The repeated emergence of the same mutations across time and lineages suggests the action of positive selection and a functional role for the less-characterized mutations. Nevertheless, the reasons underlying the genetic success of these mutations and the role of the genetic background remains to be defined. A spike mutation common to almost all the strains circulating today is S:D614G. This mutation appeared early in the pandemic and has been associated with higher fitness in experimental work *in vitro*, *in vivo* and in epidemiological settings (Korber et al. 2020; Plante et al. 2021; Yang et al. 2021). In particular, S:D614G has been reported to affect the conformational landscape of the spike glycoprotein (Yurkovetskiy et al. 2020; Benton et al. 2021; Zhang, J. et al. 2021; Gobeil et al. 2021), so that a particular domain known as the Receptor Binding Domain (RBD), discussed further in this work, may be more exposed for increased interactions with the main cellular receptor ACE2 (Ozono et al. 2021). As a consequence, S:D614G variants outcompeted the initial 614D Wuhan-Hu-1 variant across the globe, although the S:D614G mutation alone cannot explain the success of the associated variants and probably there was a combined effect with a founder event (van Dorp et al. 2020). Likewise, a new variant emerged during the summer of 2020, 20E/EU1, which was characterized by the additional spike mutation, S:A222V. This mutation has been seen both before and after summer 2020 in other genetic backgrounds, including Delta (as indicated in the **Results** section), suggesting that the mutation had a functional effect and may explain the success of the associated variant. However, in-depth analysis of the population dynamics of the variant in different countries suggested that the success of 20E/EU1 was a by-product of lifting restrictions in Europe after the first lockdown and holiday-associated travels across Europe (Hodcroft et al. 2021). This opens the question of why S:A222V has been selected multiple times in the evolution of SARS-CoV-2 and what the functional role of the mutation in viral replication or transmission is.

Here, we use S:A222V as an example of those mutations that have arisen multiple times in the SARS-CoV-2 spike protein in different genetic backgrounds but had no associated increased epidemiological fitness. Tracing the interactions with the genetic background in which they appear (epistasis) could help in predicting the potential success of these new mutations in the population. To clarify the role of S:A222V, we have used an extensive genomic dataset describing their population dynamics across SARS-CoV-2 lineages. We have also studied the role of the genetic background by combining the mutation S:A222V with S:D614G in neutralization experiments with pseudotyped viruses, followed by biophysical characterization of the mutant spike. On the structural side, we report here the cryo Electron Microscopy (cryo-EM) map and structural model of the 1-up conformation of 20E/EU1 (7QDG, EMD-13916) and the ubiquitous S:D614G mutant (7QDH, EMD-13919). Cryo-EM and Molecular Dynamics (MD) simulations were finally combined to obtain indications about the functional role of the mutation. This multidisciplinary approach allowed us to provide a plausible answer to the repeated emergence of the S:A222V mutations, which could be summarized as an allosteric effect resulting in an enhanced flexibility of the RBD in the up state, although its translation into significant phenotypic effects depends on the genetic background. Finally, serological neutralization studies and *in vitro* thermostability assays did not show a marked difference between the two strains, suggesting it does not affect vaccine effectiveness.

## Results

### Population dynamics of S:A222V through space and time

The mutation S:A222V first arose within Lineage B.1.177, described for the first time in Spain in Summer 2020 (Hodcroft et al. 2021). S:A222V has been observed in 290,789 sequences of 123 countries by September 2021, almost exclusively co-occurring with S:D614G (99.86%). Additionally, 9% of 3,227,996 SARS-CoV-2 sequences available until 2021-10-24 include [S:A222V + S:D614G] and only 0.012% harbour S:A222V without S:D614G, indicating that the presence of S:A222V is tightly linked to S:D614G. Interestingly, although S:A222V arose with B.1.177, by September 2021 more than half of the sequences with S:A222V were not in the genomic context of B.1.177, indicating that although B.1.177 was replaced by other variants (Hodcroft et al. 2021), S:A222V emerged independently in different lineages. Non-B.1.177 sequences with S:A222V include those isolated in 123 countries and classified in as many as 192 PANGO lineages, which indicates subsequent and independent appearances of S:A222V in different genetic backgrounds. By October 2021, the majority of the non-B.1.177 occurrences appear in lineage B.1.617.2, where 10% of B.1.617.2 contain S:A222V. B.1.617.2 is also known as Variant Of Concern (VOC) Delta and it includes all AY lineages (ref: https://www.who.int/en/activities/tracking-SARS-CoV-2-variants). Less frequently, S:A222Vappears in 0.11% of B.1.1.7 (VOC Alpha, ref: https://www.who.int/en/activities/tracking-SARS-CoV-2-variants), and 0.56% of B.1 overall. The temporal pattern for S:A222V (pink line in **Figure 1a**) shows two peaks, the first corresponding to the dynamics of 20E (green line in **Figure 1a**) and the second to the expansion of B.1.617.2 (VOC Delta) (yellow line in **Figure 1a**). We did not observe a similar peak in S:A222V during the increase of B.1.1.7 (VOC Alpha; blue line in **Figure 1a**), possibly suggesting that epistatic interactions may favour the transmission of S:A222V preferentially in some genetic backgrounds.

**Figure 1.**
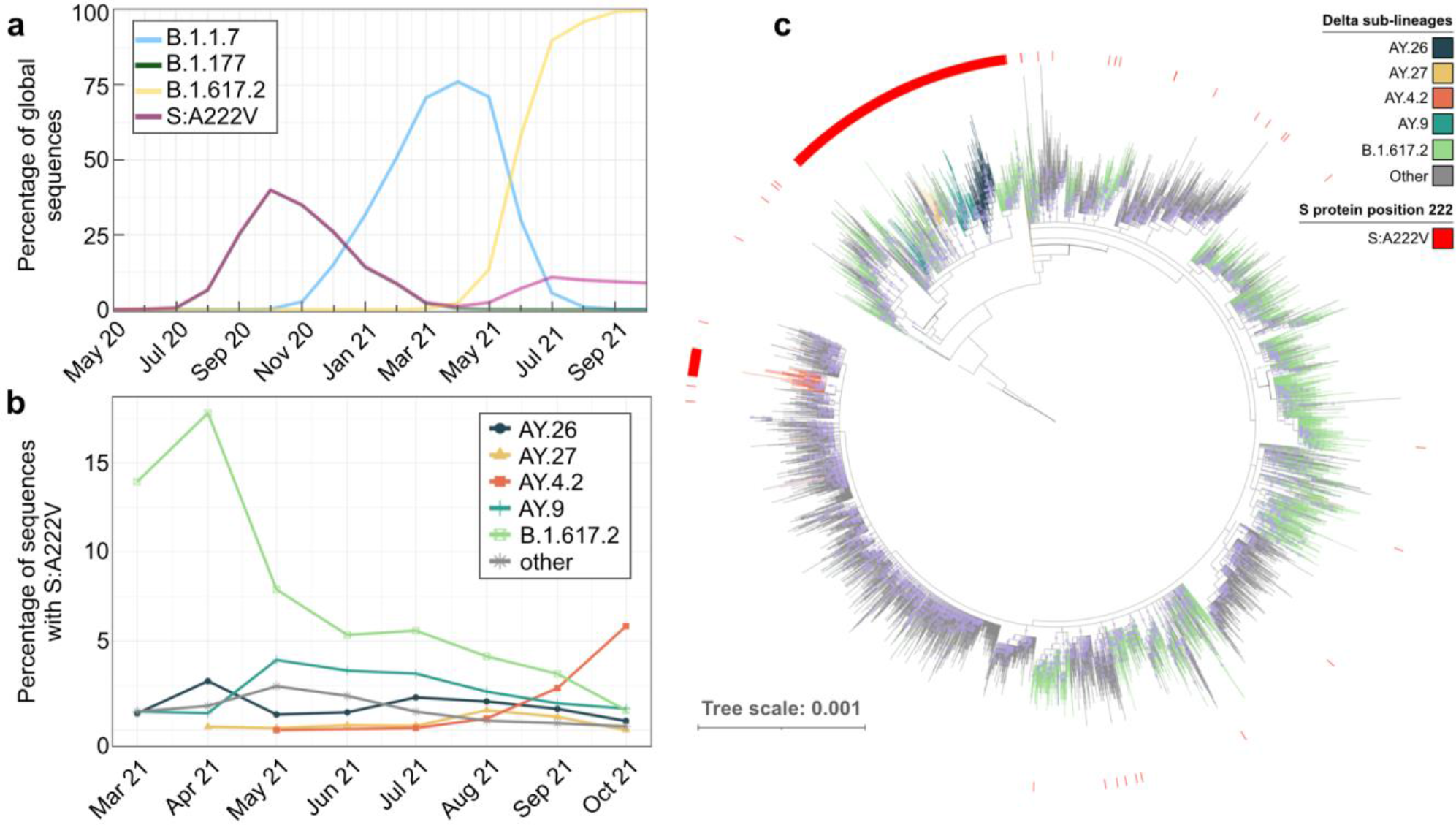
Temporal distribution of the S:A222V sequences. (**a)** Percentage of global sequences with S:A222V (pink), and sequences belonging to parental lineages designated as B.1.177 (green), B.1.1.7 (blue) and B.1.617.2 (yellow) by month. (**b)** Percentage of sequences designated as parental B.1.617.2 and its sub-lineages (designated as AY) with S:A222V by month. (**c)** Maximum likelihood phylogeny of 10,241 sequences rooted with reference MN908947.3 from Lineages B.1.617.2 and derivatives. Red external ring indicates sequences with S:A222V, the colour of branches indicates B.1.617.2 sub-lineages of interest. Each circle in branches correlates with the bootstrap value; only bootstraps from 70 to 100 are represented.

To decipher the recent dynamics of S:A222V, we focused on B.1.617.2 in a global dataset of 1,453,295 sequences (**Figure 1b** and **c**). B.1.617.2 sequences with S:A222V belong to different sub-lineages, being characteristic of four sub-lineages: AY.4.2, AY.9, AY.26 and AY.27, where it is present in more than 95% of the sequences of each sub-lineage (**Figure 1c**). However, S:A222V is also present in 25% of sequences classified as the parental B.1.617.2 lineage, and in 1.31% of other B.1.617.2 sub-lineages (**Figure 1c**). S:A222V has appeared at least 32 times in the genomic context of B.1.617.2, being detected in at least 176 different sub-lineages (**Figure 1c**).

S:A222V appeared within B.1.617.2 as early as March 2021, when it accounted for 17% of all B.1.617.2 sequences (**Figure 1b)**. After that, B.1.617.2 sequences with S:A222V increased to 23% in April 2021 but dropped afterwards. This decrease is mainly driven by the decrease of the sequences designated as parental lineage B.1.617.2 (**Figure 1c** and **Supplementary Figure 1**). Conversely, since July 2021, the proportion of B.1.617.2 (and S:A222V along with it) has been increasing mainly due to the increase of AY.4.2 (**Figure 1b)**. Among the different appearances of S:A222V in the different genomic backgrounds, it has been successfully transmitted on at least two occasions: one in the last common ancestor of AY.4.2 and the other in the common ancestor of Lineages AY.9, AY.26 and AY.27 (**Figure 1c**). Additionally, eight other less abundant lineages which derive from the same common ancestor as AY.9, A.26 and AY.27 also have S:A222V (**Figure 1c**).

### Neutralization of pseudotyped VSV by convalescent sera

One possibility for the selection of S:A222V could be an improved ability to replicate in immune populations. As vaccination had not yet started in Spain when this mutation was first observed in 2020, any such effect would have to result from selection for escape from existing immunity in convalescent individuals. To test this possibility, we evaluated the ability of VSV pseudotyped with either the ancestral Wuhan-Hu-1 spike protein, S:D614G, or 20E ([S:A222V + S:D614G]) to be neutralized by sera from the first wave of infection in Spain (April 2020) when no S:A222V mutations were circulating (**Figure 1**). No significant differences were observed in neutralization between the different viruses (**Figure 2)**.

**Figure 2.**
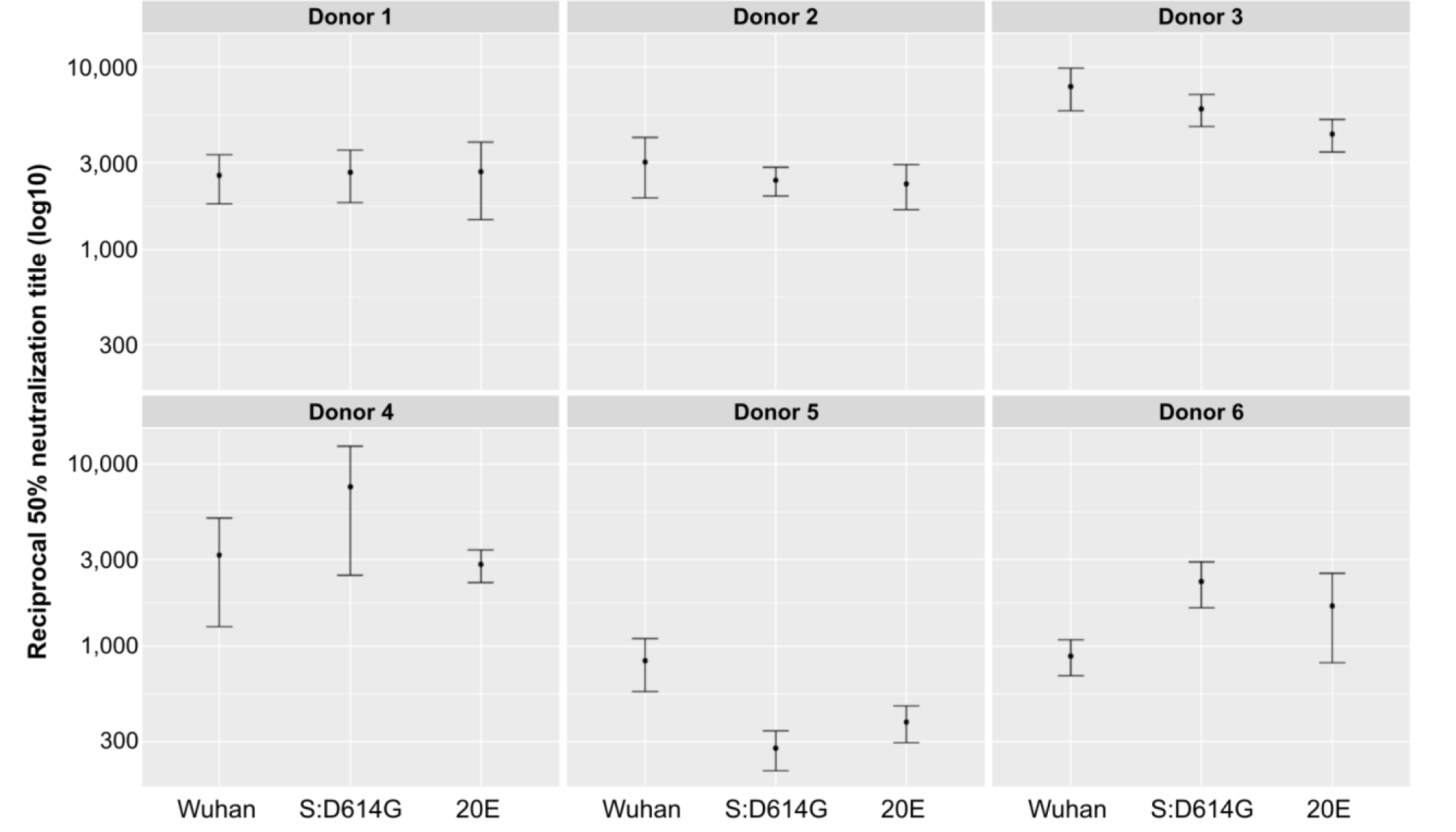
S:A222V does not alter the neutralization capacity of convalescent sera. The sensitivity to neutralization by convalescent sera of pseudotyped VSV carrying the ancestral Wuhan-Hu-1 (Wuhan), S:D614G, or the [S:A222V + S:D614G] (20E) spike protein was evaluated. No significant differences were detected between any of the sera (n=6; p>0.05 by Kruskal-Wallis test; p>0.05 by Mann-Whitney test for Wuhan-Hu-1 vs. S:D614G, Wuhan-Hu-1 vs. 20E, or S:D614G vs. Wuhan-Hu-1).

### *In vitro* functional assays

The ectodomain of SARS-CoV-2 protein S corresponding to the Wuhan-Hu-1, S:D614G or [S:A222V + S:D614G] variants, hosting proline substitutions at residues 986 and 987, a mutant non-cleavable sequence at the furin site and a C-terminal foldon trimerization motif, were flash-purified (1 day) identically and used fresh. The three forms were indistinguishable by SDS-PAGE and eluted identically from a SEC column (**Supplementary Figure 2a-b**), as expected if they had essentially the same size and glycosylation pattern. Their quality, assessed by negative-stain electron microscopy (EM), was also similarly high (**Supplementary Figure 2c**).

To functionally characterize the purified protein S variants, we first used thermal shift assays to measure their thermostability and we found that [S:A222V + S:D614G] and S:D614G variants show very similar half-melting temperatures (T0.5) and are both slightly more stable than the original Wuhan-Hu-1 variant (**Table 1** and **Supplementary Figure 3**). Then, to see the possible impact of the S:A222V mutation on the spike binding capacity to the protein ACE2, we carried out protein-protein interaction assays by biolayer interferometry (BLI) (**Table 1** and **Supplementary Figure 4**). Their affinities for ACE2 were somewhat higher for the [S:A222V + S:D614G] and S:D614G variants than for the Wuhan-Hu-1 variant (KD values of 50 nM, 66 nM and 79 nM, respectively), in agreement with previous reports comparing the S:D614G and Wuhan-Hu-1 variants (Ozono et al. 2021). Although the dissociation constants for proteins carrying S:D614G and [S:A222V + S:D614G] mutations are similar, we observed a higher kon for the [S:A222V + S:D614G] mutant, perhaps reflecting structural differences related to higher accessibility of the RBD in this mutant compared to that in the S:D614G or the Wuhan-Hu-1 variants. To investigate this possibility further, we determined the cryo-EM structures of both the [S:A222V + S:D614G] and the S:D614G S proteins.

**Table 1.**
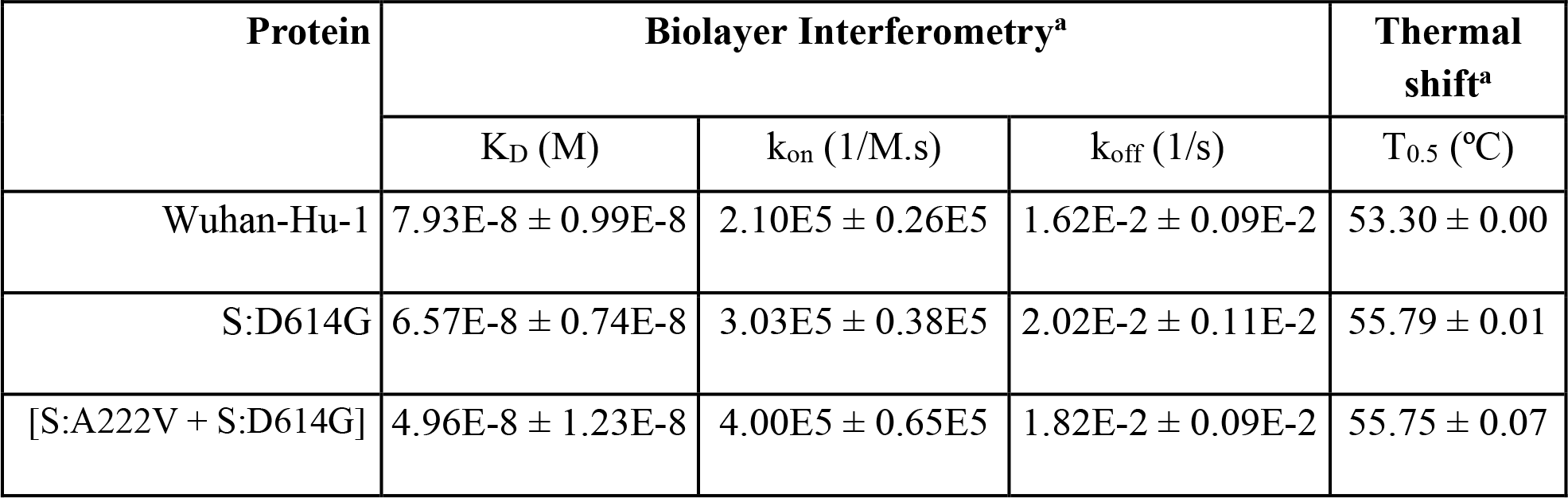
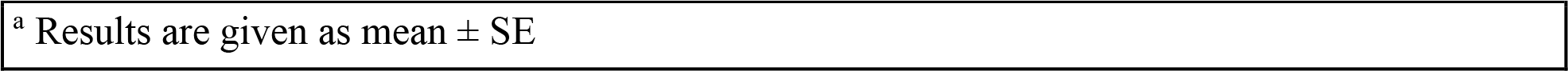
In vitro functional assays.

| Protein | Biolayer Interferometry<sup>a</sup> |  |  | Thermal shift<sup>a</sup> |
| --- | --- | --- | --- | --- |
| | $K_D$ (M) | $k_{on}$ (1/M.s) | $k_{off}$ (1/s) | $T_{0.5}$ (°C) |
| Wuhan-Hu-1 | $7.93E-8 \pm 0.99E-8$ | $2.10E5 \pm 0.26E5$ | $1.62E-2 \pm 0.09E-2$ | $53.30 \pm 0.00$ |
| S:D614G | $6.57E-8 \pm 0.74E-8$ | $3.03E5 \pm 0.38E5$ | $2.02E-2 \pm 0.11E-2$ | $55.79 \pm 0.01$ |
| [S:A222V + S:D614G] | $4.96E-8 \pm 1.23E-8$ | $4.00E5 \pm 0.65E5$ | $1.82E-2 \pm 0.09E-2$ | $55.75 \pm 0.07$ |

### Structural analysis of the allosteric role of S:A222V and epistatic interactions with S:D614G

To provide a comparative structural analysis of the S:A222V mutation on the background of S:D614G, we combined cryo-EM and classical MD simulations. Samples of spikes with both [S:A222V + S:D614G] and S:D614G, as an internal control, were produced for cryo-EM as described in **Methods**. Data about cryo-EM on the S:D614G mutant as well as a detailed description of the data sources, the modelled systems, the MD protocols and structural stability is reported in **Methods** and **Supplementary Information**.

**Cryo-EM of the [S:A222V + S:D614G] mutant.** Movies were collected as described in **Methods**. A representative micrograph is shown in **Supplementary Figure 5**. We followed both a “standard” Single Particle Analysis workflow, shown in **Supplementary Figure 6a**, and a modified one specially aimed at detecting deviation from axial symmetry through symmetry relaxation, presented in **Supplementary Figure 6b**. Note that all 3D classifications have been performed multiple times (Sorzano et al. 2021) and that assignment into classes is provided with mean and standard deviation information.

The sample, under the conditions indicated in **Methods**, presented a majority conformation 1-up with about 50% of the particles (after consensus), although two minority but stable 2-up (1.6%) and 3-down (1.9%) conformations were also found through symmetry relaxation (**Supplementary Figure 6**). The highest resolution structure of the consensus 1-up type is shown in **Figure 3a**, together with quality figures in **Figure 3b** such as the global FSC curve, the reported resolution (3.4 Å) and its angular coverage. A structural model of the 1-up conformation was derived from the map by model building and refinement procedures (**Methods** and **Figure 3c**). All structures have been submitted to the PDB and EMDB (for more details see **Data Availability**). As a control, a cryo microscopy analysis of the S:D614G mutant was also performed and presented in **Supplementary Figures 7** and **8.**

**Figure 3.**
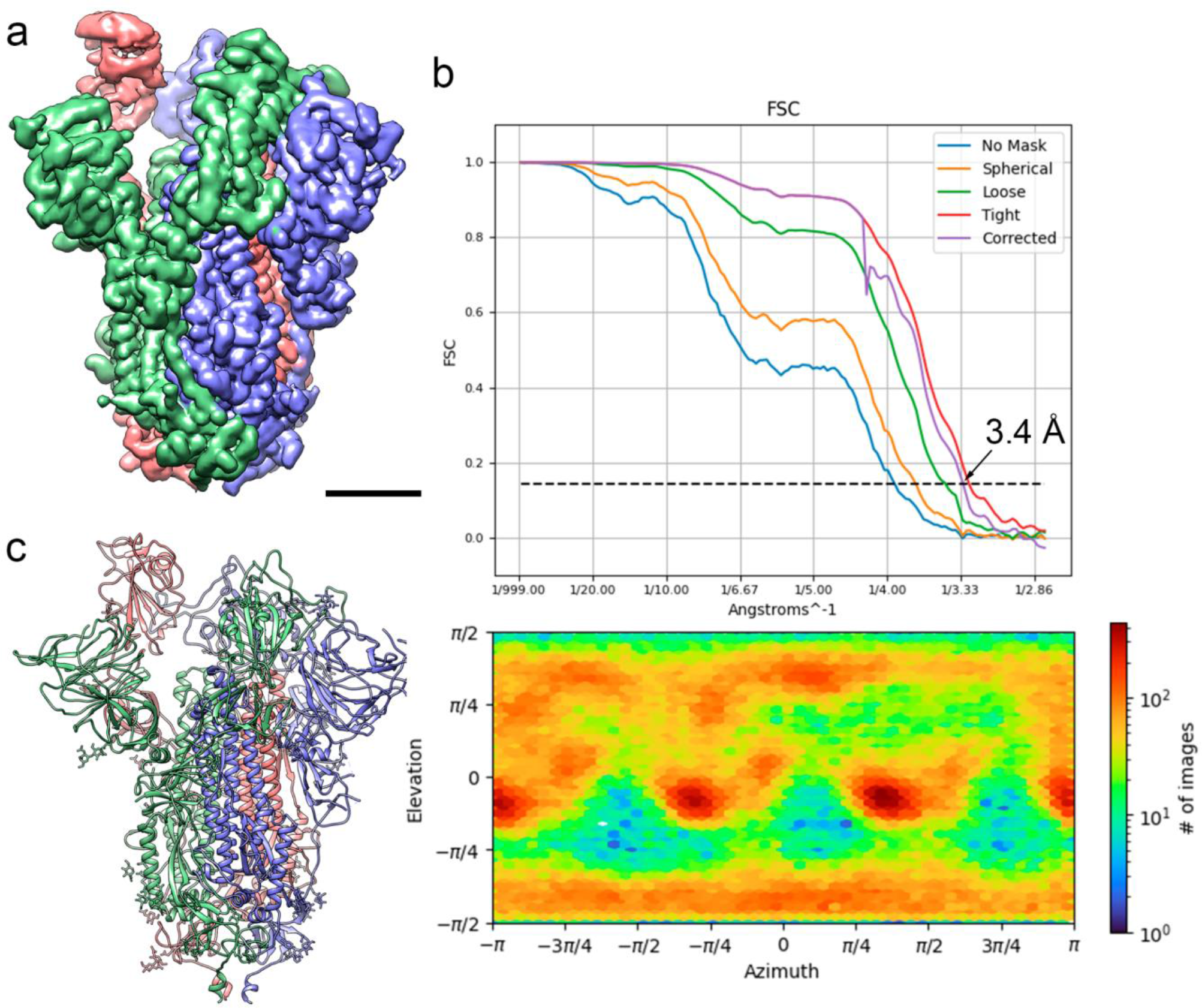
Cryo-EM of the [S:A222V + S:D614G] mutant. (**a**) Side view of the cryo-EM density map of the [S:A222V + S:D614G] mutant obtained after consensus of three independent runs of symmetry relaxation 3D classification, followed by initial model generation, refinement without symmetry imposition and deepEMhancer map sharpening, shown at 3 σ. Bar = 30 Å. (**b**) Fourier Shell Correlation (FSC) resolution curve (top) shown as the regular cryoSPARC global FSC resolution output, which includes no mask and different masks. Resolution based on the gold standard 0.143 criterion is 3.4 Å. The angular distribution coverage profile is also shown (bottom). (**c**) Atomic model of the [S:A222V + S:D614G] mutant shown as ribbon diagrams. S protein subunits are coloured in blue (chain A), red (chain B) and green (chain C). Glycan molecules are shown as stick diagrams and coloured according to their corresponding subunits.

### Structural comparison of the [S:A222V + S:D614G] and S:D614G 1-up mutants

Comparison with the structure of the S:D614G mutant reveals that in the S:A222V mutation the V222 residue is accommodated in a highly hydrophobic environment of the NTD shaped by V36, Y38, F220 and I285 (**Figure 4a**). As observed in **Supplementary Figure 9,** this mutation is expected to slightly affect the interaction of the open subunit (chain B; NTDB) with the neighbouring CTD1A. Overall, the structures for the two mutants are strikingly similar (RMSD of 1.332 Å over 3193 Cα atoms), especially at the level of the S2 region, although we can observe small conformational changes in the position of the NTDs and RBDs. These differences are particularly relevant for the RBD in position up (RBDB), which is involved in the interaction with the host receptor, ACE2. Indeed, superimposition of these two structures with that of a spike in complex with bound ACE2 (PDB: 7DF4) indicates that in these two spike structures small rearrangements of this RBD-up would be beneficial to bind ACE2 (Xu et al. 2021). The conformation of subunit C is quite similar in the two structures (RMSD 1.108 Å over 1074 Cα atoms), in contrast with the other two subunits (RMSD of 1.381 Åover 1070 Cα atoms for chain A and 1.389 Å over 1049 Cα atoms for chain B), which differ especially in (i) RBDB, (ii) RBDA, and (iii) NTDB (**Figure 4b** and **Movie 1**). The analysis of these 3 domains allows us to describe the main conformational changes between these two spike structures. RBDA is sandwiched between RBDB and NTDB. We observed that RBDA and NTDB move together as a rigid body (RMSD of these two domains between [S:A222V + S:D614G] and S:D614G structures is only of 0.726 Å), approaching RBDB in the case of [S:A222V + S:D614G] when compared to that in the S:D614G structure. On the other hand, RBDB of [S:A222V + S:D614G] undergoes a rotational movement that also brings it closer to RBDA (**Figures 4b** and **5c,** and **Movie 1**). All these movements affect the degree of tightening of the S1 region of the spike (**Movie 1**), so that the global conformation of the [S:A222V + S:D614G] mutant seems to be tighter than that of S:D614G. This is mainly a consequence of the tightening of subunit B around RBDA. In this direction, the distance between residues 114 and 381 (**Figure 4b**), located in NTDB and RBDB at the interface between these domains and the RBDA, are 3.5 Å closer in the [S:A222V + S:D614G] structure (35.3 Å in [S:A222V + S:D614G] vs. 38.8 Å in S:D614G). Although to a lesser extent, the distance between these residues in subunit A is also shorter in [S:A222V + S:D614G] when compared to that in S:D614G (42.4 Å in [S:A222V + S:D614G] vs. 43.9 Å in S:D614G).

**Figure 4.**
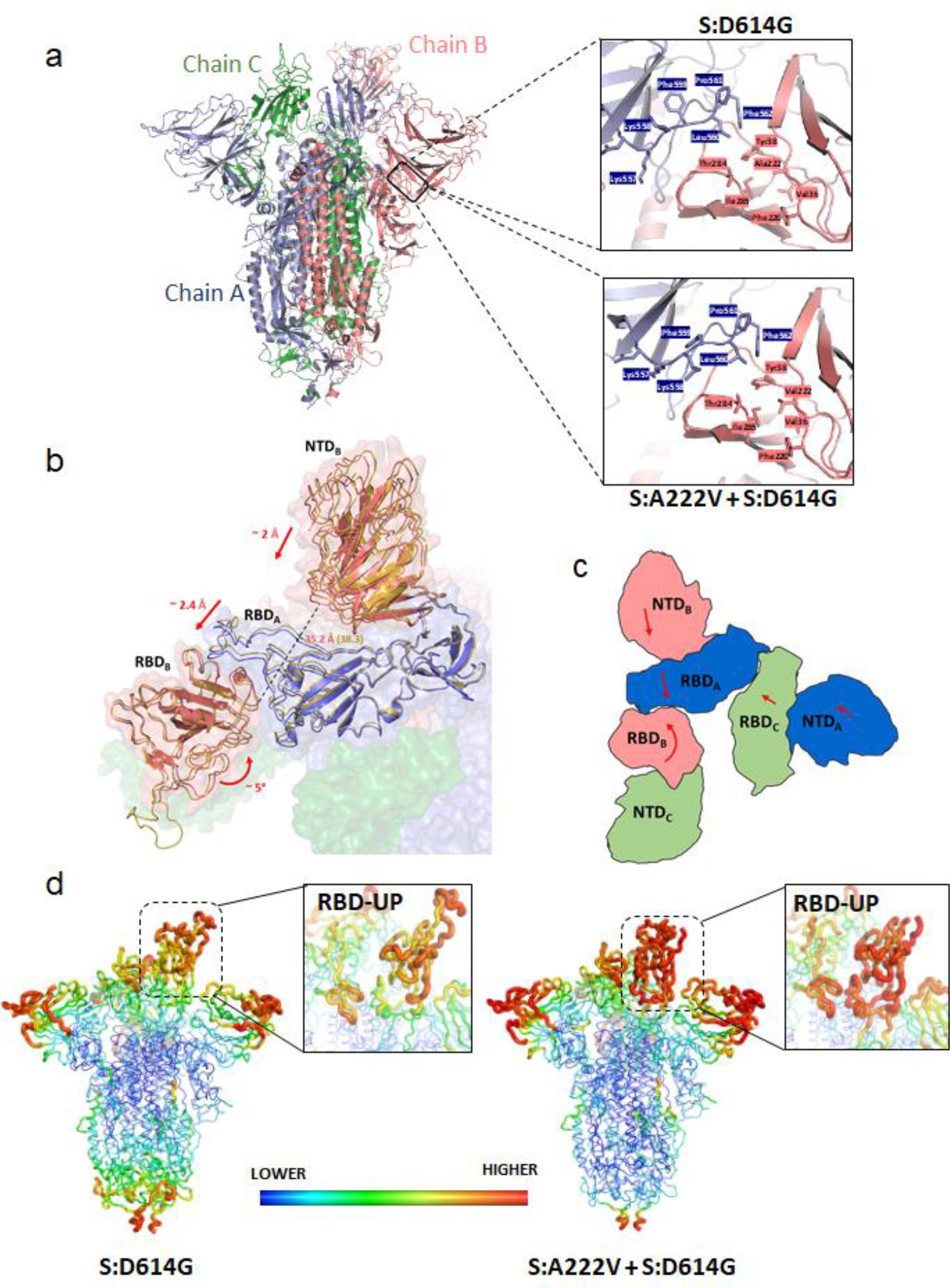
Effects of the S:A222V mutation on the structure of SARS-CoV-2 spike. (**a**) Comparison of the structures for the S:D614G and [S:A222V + S:D614G] mutants showing main conformational changes observed when S:A222V is present. Detailed view of the trimeric spike surface shown in semi-transparent representation with different colours for the different subunits, and RBDB, RBDA and NTDB domains from the S:D614G and [S:A222V + S:D614G] mutant structures shown as cartoon. Domains corresponding to the S:D614G structure are coloured in yellow. Arrows indicate the direction and magnitude of the observed domain movements. A dashed line represents the distance between the Cα atoms of residues 114 and 381 in chain B, which is ∼ 3 Å higher in S:D614G than in [S:A222V + S:D614G]. (**b**) Schematic representation of the RBD and NTD from different subunits to show the movements observed when comparing the two structures (**c**) On the left, cartoon representation of the trimeric spike. On the right, detail of the region of interaction between the NTD from subunit B (salmon) and the CTD1 region from subunit A (blue). The side chains of residues surrounding A222 (upper panel) or V222 (lower panel) are shown as sticks. (**d**) Comparison of the B-factor of RBDB of the two spike mutants. Backbone is coloured and sized according to the B-factor values for S:D614G (left) and [S:A222V + S:D614G] (right).

**Figure 5.**
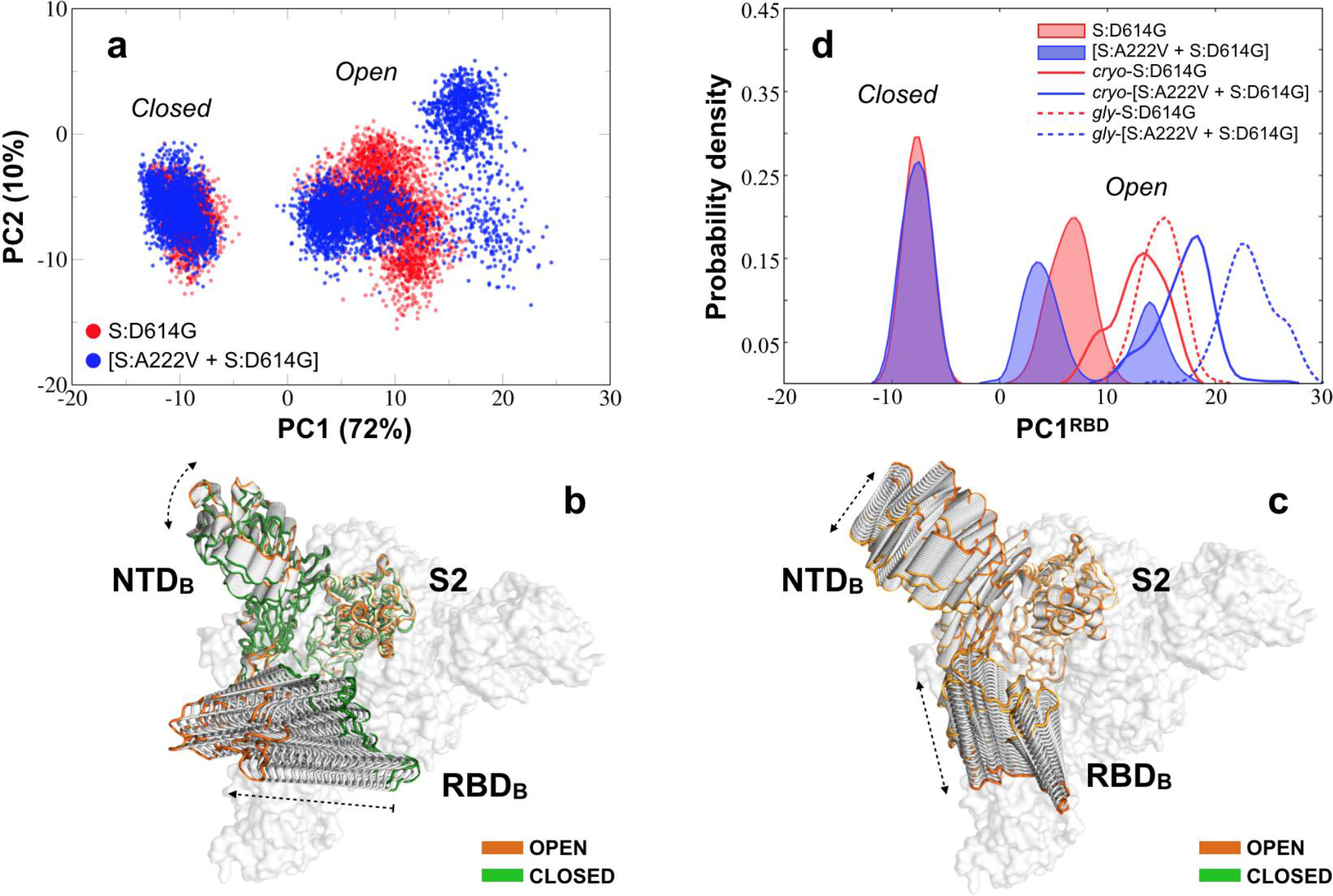
PCA for the three MD simulated systems. (**a**) PCA projection of the first two high-variance motions for the S:D614G (in red) and [S:A222V + S:D614G] (in blue) mutants of SARS-CoV-2 S1 subunit of the glycan-free spike protein. (**b)** Conformational evolution of the NTD and RBD along the first eigenvector (PC1) mainly accounting for the RBD *closed-to-open* transition. The open and closed states at the two ends of the PC1 are highlighted in orange and green, respectively. (**c**) Conformational evolution of the NTD and RBD along the second eigenvector (PC2) accounting for lateral RBD+NTD fluctuations. An orange gradient is applied here to distinguish between different lateral orientations of the open NTD-RBD pair at its two ends. (**d**) Continuous population densities along the PC1 for a PCA relative to the RBD (residues 330-530; PC1^RBD^): solid profiles are used for the simulated closed, 3-down and open, 1-up states from the glycan-free systems based on 6VXX and 6VSB while continue/dashed lines are used to define population densities for the simulated open 1-up states of the cryo-EM/fully-glycosylated S:D614G (red) and [S:A222V + S:D614G] (blue) mutants.

These tightening movements also increase the surface of contact between the NTD and RBD of the different subunits in [S:A222V + S:D614G] when compared to those in S:D614G (**Supplementary Table 1**). This effect seems to correlate with neither an increase in the stability of the S protein, as suggested by thermal shift stability assays (see above), nor with a decrease in the flexibility of the protein. In fact, an analysis of the temperature factors (B-factor) of the two structures (**Figure 4d**) shows a much higher mobility for RBDB in [S:A222V + S:D614G]. Not surprisingly, the regions in direct contact with RBDB (that is, RBDA and the contacting loops of NTDC) also show increased B-factor values.

### Molecular dynamics simulations of the S:D614G and [S:A222V + S:D614G] mutants

300 ns MD simulations run in triplicate for each of the two simulated systems in the 3-down (DDD), closed state, yielded a total of 0.9 μs of sampling per system. Given the metastable nature of the open, 1-up (UDD) conformation of the Spike protein, longer 500-ns simulations were run in this case, totalling 1.5 μs per system (DDDS:D614G, DDD[S:A222V + S:D614G], UDDS:D614G, and UDD[S:A222V + S:D614G] models in **Supplementary Table 2**). 200 ns of MD simulations were also run for both S:D614G and [S:A222V + S:D614G] mutants based on atomistic structures fitted to our cryo-EM densities (*cryo*-UDDS:D614G and *cryo*-UDD[S:A222V + S:D614G] models in **Supplementary Table 2**).

We initially analysed these simulations *via* principal component analysis (PCA) focusing on a single S1 subunit (**Methods**). The projections on the first two eigenvectors for the MD simulated mutants, collectively capturing 82% of the structural variance within the conformational landscape explored by the S1, are shown in **Figure 5a**. The largest motion represented by the first eigenvector (PC1: 72%) mainly describes the fluctuation of the RBD from the closed to the open state, with a minor contribution from the NTD of the same subunit (**Figure 5b**). The second motion (PC2: 10%) represents the lateral fluctuation of the RBD along with lateral movements of NTD (**Figure 5c**), which affects the orientation of the solvent-exposed RBD region in the open state. As shown in **Figure 5a**, while the closed states of both mutants populate an overlapping well-defined region of the PCA space, the open state of the [S:A222V + S:D614G] mutant is observed to explore a visibly larger range of conformations in a clearly multimodal manner.

To more directly trace the *closed-to-open* transition of the RBD, we recalculated the PCA using RBD atoms (residues 330-530) and plotted continuous population densities along PC1 for the open and closed RBD domain of the 1-up and 3-down models of the two mutants (**Figure 5d**). As expected from **Figure 5a**, the open RBD of the [S:A222V + S:D614G] mutant sampled a bimodal density distribution (values centered on 4 and 14 arbitrary units for PC1^RBD^), with the leftmost substate largely overlapping with the unimodal distribution of the S:D614G mutant (values centered on 8 for PC1^RBD^; **Figure 5d**). Density distributions for the S:D614G and [S:A222V + S:D614G] mutants from the simulations of the 1-up cryo-EM structures are also reported in **Figure 5d** (unfilled solid lines). Comparison with the reference distributions (red and blue filled profiles in **Figure 5d**) revealed that both are slightly shifted to more positive values (respectively centred on 14 and 19) along the PC1^RBD^.

Let us remark that the cryo-EM [S:A222V + S:D614G] mutant still explores a more open distribution with respect to the S:D614G mutant, thus confirming our finding. Likewise, it seems to have no observable effect on the closed 3-down state.

To characterize the relative impact of glycosylation on the conformational preferences observed for the previously analysed glycan-free systems, we also simulated both mutants from the fully glycosylated structure (**Methods**). Since no relevant effects, relative to the observed phenomenon, were produced by the S:A222V mutation on the conformational ensemble of the 3-down closed state (see above), MD simulations were limited to the open 1-up state of the protein for this analysis. Accordingly, 400 ns of MD simulations were run for the S:D614G and [S:A222V + S:D614G] mutants (*gly*-UDDS:D614G and *gly*-UDD[S:A222V + S:D614G] models in **Supplementary Table 2**). As before, the *closed-to-open* transition of the RBD was analysed by performing PCA on the RBD (residues 330-530) and continuous density distributions for PC1^RBD^ were finally plotted (red and blue dashed lines in **Figure 5d**). In line with other published data (Casalino et al. 2020; Pang et al. 2021; Rahnama et al. 2021), glycosylation seems to affect the extent of the RBD opening. Even more shifted density distributions were in fact registered for both the S:D614G and [S:A222V + S:D614G] mutants (respectively centred on 15 and 22 for PC1^RBD^) with respect to the previously simulated systems. Nevertheless, we still observe the stabilization of a more open conformation of the RBD for the glycosylated [S:A222V + S:D614G] mutant, thus confirming the general trend registered for our more extensive, microsecond-scale explorations relative to the glycan-free mutants (**Figure 5a** and **5d**).

To account for potential interdomain allosteric implications, we also analysed the effects induced by the RBD opening on the other two closed RBDs within the same trimeric ensemble of the simulated 1-up trajectories (**Supplementary Figure 10a, c,** and **e**). For completeness, data obtained from this analysis were also compared with those for the whole S1 units (**Supplementary Figure 10b, d,** and **f**). A slightly higher dynamical behaviour was observed at the level of the closed S1/RBD units for the simulated (glycan-free) cryo-EM mutants (**Supplementary Figure 10c** and **d**), with values spreading between -15 and 5 along the PC1^S1/RBD^. A tendency to more negative values for the two closed subunits (blue vs. red profiles in **Supplementary Figure 10c** and **d**) of the [S:A222V + S:D614G] mutant was observed, especially for the glycan-free cryo-EM trajectories, thus suggesting the possibility of a certain uncoupling of its trimeric S1 (RBD-NTD) ensemble. We also performed a mutational free energy analysis on the open, 1-up structures (**Methods**). Results from this analysis (**Supplementary Table 3**) indicated that the impact of the S:A222V mutation on the RBD opening is too small to establish a significant preference for either of the conformational states of the RBD.

### Dynamic network analysis

To identify direct mechanical connections responsible for a more open RBD conformation in the [S:A222V + S:D614G] mutant, we performed dynamic network analysis on an adjacent RBD-NTD pair (see **Figure 6a** and **b**), i.e. with the two domains coming from two different chains.

**Figure 6.**
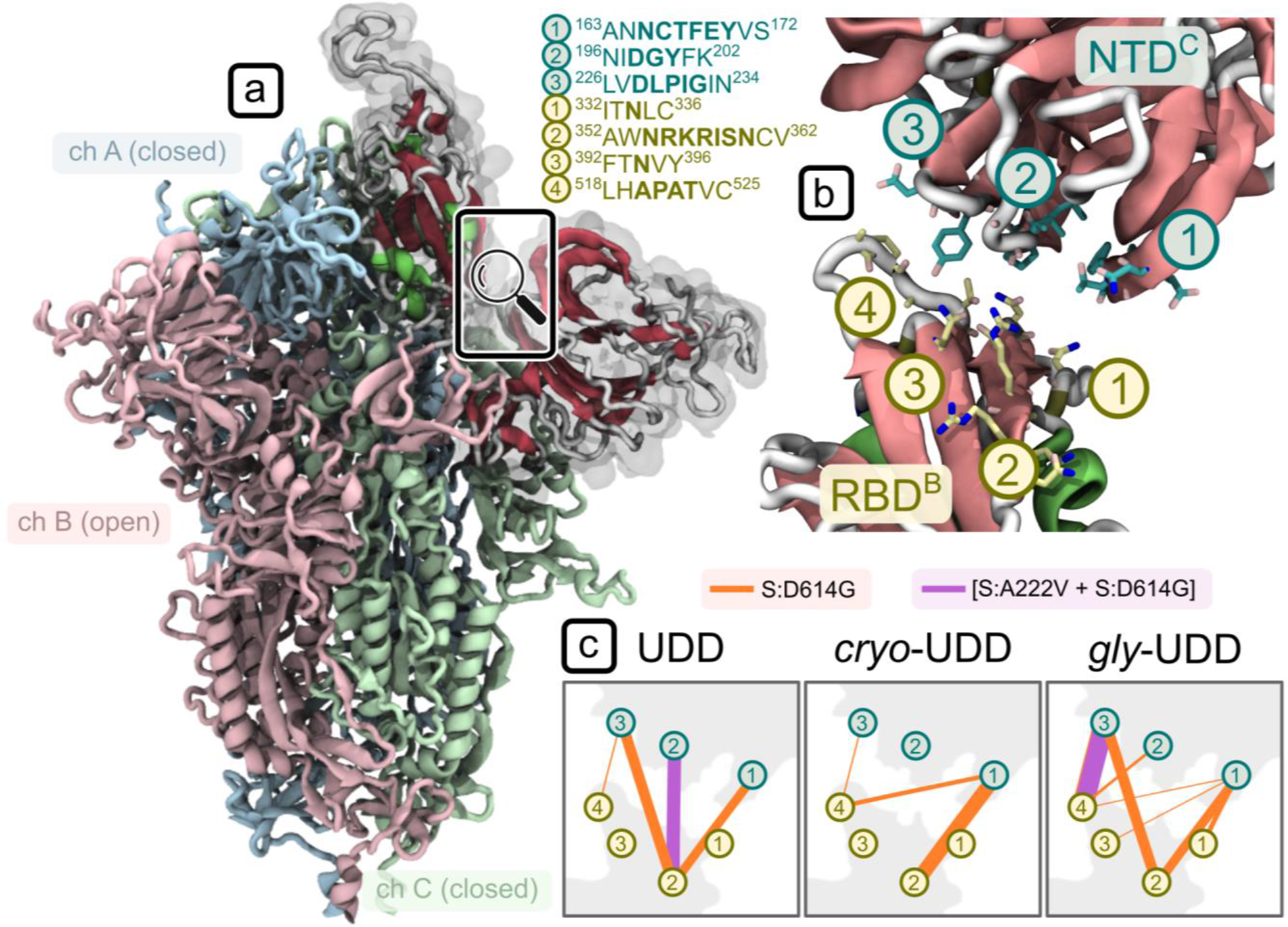
Dynamic network analysis for a subsystem consisting of the open RBD and its neighbouring NTD. (**a**) Open RBD of the simulated 1-up (UDD) SARS-CoV-2 Spike mutants (shaded in A). (**b**) Residue communities (3 for NTD, in teal, and 4 for RBD, in yellow) engaged in inter-domain contacts, along with their position in the protein sequence. (**c**) Communication pathways are highlighted with connecting lines whose thickness is proportional to the inter-domain betweenness centrality values. Orange connections represent the S:D614G and violet the [S:A222V + S:D614G] mutant; data is reported for the three systems: glycan-free from 6VSB (UDD), glycan-free from our cryo-EM data (*cryo*-UDD), and glycosylated from 6VSB (*gly*-UDD).

The relative values of betweenness centrality, reported in **Figure 6c**, show a dramatic difference in the extent of connectivity between the two domains which is mainly driven by hydrophobic contacts. In the three datasets, the [S:A222V + S:D614G] mutant forms at most a single strong “pivot” connection between the two domains (see violet community pairs 2^NTD^-2^RBD^ and 3^NTD^-4^RBD^ for UDD and *gly*-UDD respectively in **Figure 6c**), thereby allowing for a degree of flexibility in their relative geometry, while the S:D614G mutant forms 3-6 stable contacts (mainly involving communities 1 and 3 of the NTD with 2 and 4 of the RBD in **Figure 6c**). This last network is extensive enough to stabilize the RBD in a single dominant conformation. No stable connections were found between the RBD-NTD pair of the cryo-EM [S:A222V + S:D614G] mutant below the selected contact cut-off of 4.5 Å. As above, this trend shows up consistently across the three different setups - i.e., independent of both glycosylation and the source of the initial structure.

### Structural flexibility of PDB-reported S:D614G single mutant

In an attempt to provide more details about the dynamics of the S:D614G mutant, especially from a wider timescale perspective, a set of 24 structures of S:D614G previously reported in the PDB were also analysed by PCA together with our new cryo-EM structures. Note that the whole process of sample preparation for cryo-EM may take from seconds to minutes, therefore we expect that the individual images that support these PDB structures came from virtually any possible conformational state of the spike, although the maps themselves are the result of an image processing process which might not be able to accurately follow these structural variations. We also remark that for structures analyzed in this PDB-wide approach, specimens may differ among themselves in more mutations than S:D614G and S:A222V alone, related to different ways to aid purification of stable spike trimers e.g. adding tags, eliminating cleavage sites and introducing proline mutations (**Supplementary Table 4**); still, all these structures were pooled together and analysed using PCA (**Figure 7** and **Supplementary** Figures 11 and 12).

**Figure 7.**
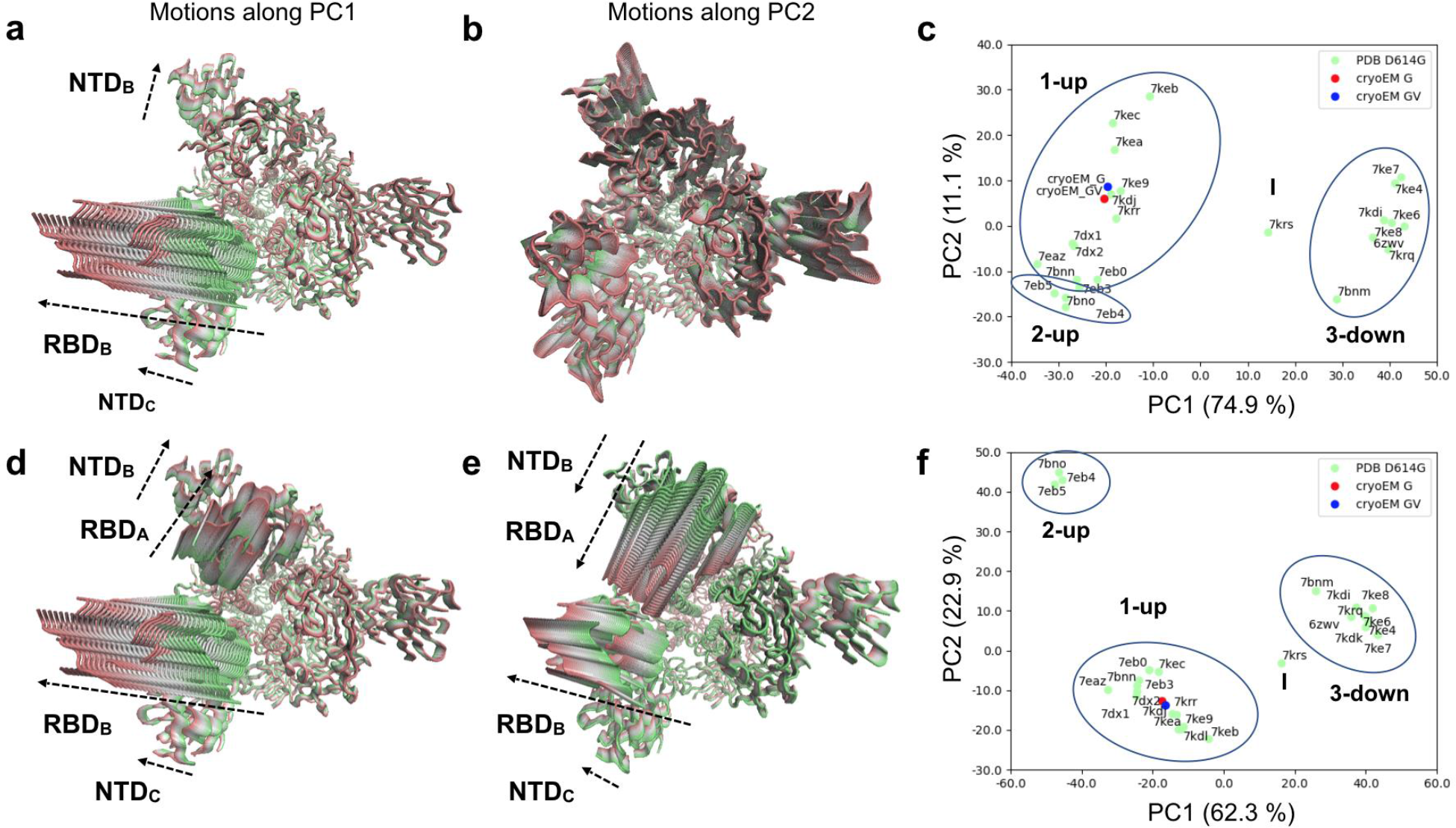
PCA for PDB-reported S:D614G single mutant. PCA results for the whole spike excluding **(a-c)** and including the 2-up conformations **(d-f)** in the covariance calculation. **(a)** Conformational evolution showing RBD opening motions observed in PC1 of the whole spike excluding 2-up conformations is shown from a top view ranging from a closed state (green) to an open state (red). **(b)** Conformational evolution along PC2 of the whole spike excluding 2-up conformations is shown from a top view from one extreme in green (compressed state) to the other in red (stretched state). Different domains move differently besides the compression and stretching of the structure. **(c)** Projection of the PDB S:D614G structures (green) and our cryo-EM structures (red and blue) onto the space of the first two PCs of the whole spike excluding 2-up conformations. Note a flipping of PC1 in this analysis relative to the PCAs from MD ensembles. **(d)** Concerted RBD opening motions observed in PC1 of the whole spike including 2-up conformations are shown from a top view ranging from one extreme in green (closed state) to the other in red (open state). **(e)** Anticorrelated RBD opening motions observed in PC2 of the whole spike including 2-up conformations are shown from a top view ranging from one extreme in green (closed state) to the other in red (open state). **(f)** Projection of the PDB S:D614G structures (green) and our 1-up cryo-EM structures (red and blue) onto the space of the first two PCs of the whole spike including 2-up conformations.

The much smaller size of the ensemble as compared to that from MD simulations allowed us to analyse the dynamics of the whole spike, which produced very similar motions of the dominant subunit to those seen when focusing on the first S1 subunit involved in RBD opening as above, especially in the first three eigenvectors (**Supplementary Figure 11a-b**), while allowing us to identify motions of NTDs and RBDs from other subunits coupled to large motions of the dominant RBD (**Figure 7a** and **Supplementary Figure 11c**). However, to maintain a direct comparison with the MD data, we initially excluded the 2-up structures from the covariance calculation to avoid the extra component of variation associated with the second RBD opening, which has not been explored in our MD simulations.

In line with the PCA from MD simulations (**Figure 5**), the first eigenvector or principal component (PC1: 89.4 %) from experimental S:D614G structures based on the single subunit is associated to the *closed-to-open* transition for the RBD together with a rearrangement of its NTD towards and away from the RBD. The equivalent motion of this subunit (chain B in our structures) in PC1 of the whole spike (overlap 0.99 and contribution 74.9 %; **Supplementary Figure 11a-b**) also featured motions of the neighbouring NTD (chain C in our structures), which moved together with the dominant RBD (**Figure 7a**). The second eigenvector of both PCAs had a low contribution (4.9% and 11%) and was dominated by stretching and compression within it (**Supplementary Figure 11d**). This non-physiological variation in size of the structure is likely related to errors in cryo-EM data acquisition, processing and/or modelling. However, there are also more potentially interesting contributions of particular domains in this mode, including the second and third RBDs and the associated NTDs (**Figure 7b**), making it interesting to include.

As with the MD, projection of these 24 structures together with our new cryo-EM structures of the S:D614G and [S:A222V + S:D614G] mutants into the space of these two PCs (**Figure 7c**) allowed a separation of 3-down and 1-up structures along PC1, as well as an intermediate structure (PDB: 7KRS) (Zhang et al. 2021). There were a number of structures with a greater degree of RBD opening relative to the two new cryo-EM structures, which occupied a similar place to each other in this landscape in line with their differences being related to more local motions, suggesting the possibility of greater opening in the S:D614G mutant too, albeit on longer timescales than accessible to the MD simulations.Interestingly, there is also some separation along PC2 including separation of some 1-up structures including one with an intermediate conformation second RBD (upper region; PDB: 7KEC) (Gobeil et al. 2021) and 2-up structures (lower region; PDB: 7BNO, 7EB4 and 7EB5) (Benton et al. 2021; Yang et al. 2021) from other 1-up structures.

Finally, to better describe the full structural variation over the longer timescale captured by this set of PDB structures, we also performed an additional PCA by including the 2-up structures in the covariance calculation, allowing us to capture the associated variation in the second RBD as well (see **Supplementary Figure 12a**). In this case, PC1 and PC2 (62% and 23% contributions; **Supplementary Figure 12b**) both showed high overlaps with PC1 of the single subunit (1.00 and 0.86, respectively; **Supplementary Figure 12c**), suggesting that they were both dominated by opening and closing of the first RBD. Intriguingly, both PCs also featured significant motions of the second RBD (chain A of our structures), which is involved in the transition to 2-up (**Supplementary Figure 12d-e**). PC1 showed coupled opening and closing of these two RBDs (**Figure 7d**); whereas PC2 showed one opening while the other closed (**Figure 7e**). There was a dominance of the first RBD in PC1 (**Supplementary Figure 12d**) and a dominance of the second RBD in PC2 (**Supplementary Figure 12e**), enabling a much clearer separation of 2-up structures in the projection onto these two PCs (**Figure 7f**). We again see that our two cryo-EM structures reside in the middle of the 1-up cluster, showing room for further opening of both dominant RBDs within the 1-up macrostate.

## Discussion

We started our study prompted by the emergence in Spain of variant 20E in summer 2020, deepening into this work as one of the key mutations characteristic of this variant, S:A222V, kept reappearing. Results from genomics, neutralization analysis, cryo-EM and MD were integrated to provide grounds to formulate a hypothesis answering some of the questions on SARS-CoV-2 evolution such as: Why do rare spike mutations keep reappearing in multiple backgrounds, including Delta, as is the case for S:A222V? It is worth mentioning that the epidemiological success of a mutation depends on the genetic background where it emerges but it can also just reflect underlying population genetic processes such as founder effects that impact the frequency of the mutation. S:A222V represents a good example of the aforementioned phenomenon. In fact, the success of 20E was likely driven by superspreading events and human mobility (Hodcroft et al. 2021) rather than S:A222V *per se*. On the contrary, the case of AY.4.2. suggests a subtle increase in transmissibility of the virus that is likely associated with S:A222V and/or S:Y145H.

Here, we have carried out an in-depth evaluation of the impact of S:A222V with the aim to capture relevant features of the mutation. This multidisciplinary approach also offered the possibility to understand how *in silico* and *in vitro* protein structure analysis can be combined to predict the impact of SARS-CoV-2 spike mutations. In this way, the biophysical characterization of two types of samples (S:D614G and [S:A222V + S:D614G]) was carried out, including thermal stability assays and protein-protein interaction assays by biolayer interferometry. These studies showed that while S:A222V does not increase the stability of the spike protein, it slightly improves its affinity towards the host receptor, ACE2. This enhanced affinity is mainly driven by an increased association constant (kon), in agreement with the increased flexibility of the RBD in the 1-up configuration suggested by both MD and cryo-EM studies.

Cryo-EM was used to obtain maps of the S:D614G and [S:A222V + S:D614G] mutants at global resolutions of respectively 4.2 and 3.4 Å from which structural models were derived. Images were processed using workflows oriented towards identifying symmetry breaking conditions characteristic of one or several of the RBDs in an up conformation. To avoid classification instabilities and inaccuracies in continuously flexible systems in which discrete flexibility classification methods were imposed (Melero et al. 2020), we performed a discrete classification analysis but enforcing results reproducibility through consensus (Sorzano et al., 2021). Under these very stringent reproducibility conditions, we were able to report a majority population of 1-up structural classes and a minority population of 2-up structural classes in both S:D614G and [S:A222V + S:D614G], with similar percentages. We only found the 3-down conformation in a very small percentage of the [S:A222V + S:D614G] mutant images. However, just indicating the percentage of particles that had been reproducibly clustered together and then visually labeled as 1-up, 2-up or 3-down is a gross oversimplification of the informational content of cryo-EM maps.

Furthermore, we might be close to the limits of what discrete 3D classification can obtain in a system like the one under study (Melero et al., 2020; Sorzano and Carazo 2021; Punjani and Fleet, 2021a). Here, a possible improvement could be provided by using new continuous flexibility approaches, whose practical limits still need to be tested (Sorzano and Carazo 2021; Chen and Ludtke 2021; Seitz et al. 2021; Punjani and Fleet 2021b; Mashayekhi et al. 2021). Nevertheless, it is important to note that refinement of the cryo-EM structures in 1-up conformation of both the S:D614G and [S:A222V + S:D614G] mutants in their corresponding maps yielded B-factors that were notably different, especially at the RBD, suggesting that this latter domain was more flexible in the [S:A222V + S:D614G] mutant. Other map-to-model quality figures, like FSC-Q (Ramírez-Aportela et al. 2021), agreed in this respect (data not shown).

Molecular dynamics simulations were also carried out to provide complementary and perhaps less discrete structural details about the subject under study. Considering the importance of glycosylation for the spike dynamics (Casalino et al. 2020; Pang et al. 2021, Rahnama et al. 2021), MD trajectories were collected starting from our cryo-EM structures as well as from previously reported structural models with and without glycosylation. Results from PCA relative to the RBD *closed-to-open* transition pointed out the preference for a more open RBD for the [S:A222V + S:D614G] mutant, with glycosylation additionally enhancing the openness of the open RBD, in agreement with a recent study (Pang et al. 2021). Furthermore, dynamic network analysis allowed us to go deeper into the allosteric nature of the possible changes induced by the S:A222V mutation. These data support the hypothesis of a more pronounced uncoupling of the RBD from its neighbouring NTD for the [S:A222V + S:D614G] mutant when compared to the S:D614G mutant alone. This effect would be due to a lower conformational restraint imposed by the NTD on RBD dynamics in the 1-up conformation as a consequence of the reduced number of connections between these two domains in the [S:A222V + S:D614G] mutant (no special effects were observed on the 3-down conformation). Of note, this trend is consistently observed across all the simulated (glycan-free and glycosylated) systems. Interestingly, a mutational free energy analysis was not able to establish a thermodynamic preference for the open conformation between S:D614G and [S:A222V + S:D614G] mutants. This would suggest that although a wider RBD opening for the [S:A222V + S:D614G] mutant is possible it wouldn’t be associated with a significant energetic gain. Finally, we compared by Principal Component Analysis all reported PDB structures of S:D614G (obtained by cryo-EM) as well as the ones presented here, revealing similar motions to those seen by MD. Clearly the structures separate into 1-up, 2-up and 3-down (and an “intermediate” one), as expected, with the ones obtained in this work essentially in the middle of the 1-up cluster. Still, further work comparing MD simulations and cryo-EM maps needs to be performed, with continuous flexibility analysis being an essential component of these future analyses.

In conclusion, the S:A222V mutation at the NTD of the SARS-CoV-2 spike sequence is able to produce small but noticeable allosteric effects in the RBD that translate into changes in its conformational dynamics as well as biophysical properties. These findings agree with the recall of the S:A222V across waves, lineages, and space, suggesting a role for positive selection. Still, the genetic background in which the mutation occurs seems to be crucial for its success and the epidemiological effects of S:A222V mutation may depend on epistatic interactions between mutations, with S:L452R and S:T478K also potentially modifying the behavior of the open RBD. Although predicting the combined outcome of different mutations on different backgrounds is still an open question, this work gets us a step closer by pinpointing the defined manner in which S:A222V affects key properties of the spike.

## Methods

### Sequence analysis

The global dataset of 3,227,996 sequences was obtained from GISAID database (Elbe and Buckland-Merrett 2017) since first case detected from 24 December 2019 until 24 of October 2021 and filtering out sequences from non-human host, with no complete collection date, with low coverage or incomplete sequences (accession ids at data table 1). 3,227,996 SARS-CoV-2 sequences were aligned with Nextalign package (https://github.com/nextstrain/nextclade/tree/master/packages/nextalign_cli). Single nucleotide polymorphisms in amino acid position 222 of the spike were obtained after generating a VCF file from each individual fasta sequence derived from the alignment using SNP sites v 2.5.1 (Page et al. 2016) with argument “-v” and using the reference genome (NC_045512.2) as the reference bases for detecting mutated sequences. B.1.617.2 sequences were obtained after selecting all sequences classified as B.1.617.2 PANGO lineage by GISAID from the global dataset described above. After eliminating sequences harbouring at least one indetermination (symbolized as “N”), and duplicated sequences detected with with seqkit v 0.13.2 (Shen et al. 2016) (arguments employed: rmdup -s) we obtained a high quality B.1.617.2 dataset of 549,382 sequences. Finally, in order to reduce the number of sequences to a suitable dataset for phylogenetic reconstruction, we selected 10,242 sequences from the high-quality B.1.617.2 dataset randomly but keeping the same temporal distribution by month, the same proportion of B.1.617.2 sub-lineage and the same proportion of sequences with amino acid V in S:222 as the initial alignment of 549,382 sequences. To determine phylogenetic relationships of B.1.617.2 and the emergence of A222V, phylogenetic analysis was performed as previously described (Ruiz-Rodriguez et al. 2021).

### Plasmids

The plasmids used for the generation of the pseudotype vesicular stomatitis virus (VSV) were previously described (Ruiz-Rodriguez et al. 2021). Plasmid pSPIKE (a generous gift from Cesar Santiago, CNB-CSIC) was designed to include the region encoding the SARS-CoV-2 protein S ectodomain (residues 15-1213) with substitutions to proline at residues 986 and 987 and of ^668^RRAR^671^ furin cleavage site to alanine, with a N-terminal gp67 signal peptide for secretion, and C-terminal foldon trimerization motif, a thrombin protease recognition site and 9x His and Myc tags, into the insect expression plasmid pFastBac. Plasmid pACE2TEV was prepared from that previously used to express the N-terminal peptidase domain of human ACE2 (Lan et al. 2020), by including a cleavage site for TEV protease, to allow removing the C-terminal 6×His tag in an additional purification step.

### Evaluation of neutralization by convalescent-phase sera

Pseudotyped VSVDG-GFP bearing the ancestral Wuhan-Hu-1, the S:D614G, or 20E (mutations S:A222V and S:D614G) spike protein were produced as previously described (Ruiz-Rodriguez et al. 2021). Neutralization assays were performed as previously described (Ruiz-Rodriguez et al. 2021). Briefly, 16 h post infection GFP signal derived from VSV replication was determined in each well using an Incucyte S3 system (Essen Biosciences). The mean GFP signal observed in several mock-infected wells was subtracted from that of all infected wells, followed by standardization of the GFP signal in each well infected with antibody-treated virus to that of wells infected with mock-treated virus. Any low or negative values resulting from background subtraction were arbitrarily assigned a low, nonzero value (0.001). The serum dilutions were then converted to their reciprocal and the dose resulting in a 50% reduction in GFP signal was calculated using the R drc package v 3.0-1. A three-parameter log-logistic regression (LL3 function) was used for all samples. All serum samples were from donors that were admitted to the intensive care unit during April 2020 (ethics approval RVB20017COVID).

### Site directed mutagenesis and protein production

Site-directed mutagenesis of pSpike was performed using Q5® Site-Directed Mutagenesis Kit (New England Biolabs) and the forward and reverse primers given in **Supplementary Table 5**. The correctness of the constructs, the presence of the desired mutation, and the absence of unwanted mutations were corroborated by sequencing.

For producing SARS-CoV-2 protein S variants (Wuhan-Hu-1, S:D614G or [S:A222V + S:D614G]) we used the Bac-to-Bac Baculovirus Expression System (Invitrogen). *E. coli* DH10Bac cells (Invitrogen), transformed with the appropiate pSpike construct carrying either Wuhan-Hu-1, S:D614G or [S:A222V + S:D614G] variants, were grown on LB-agar containing 50, 7, 10, 40 and 100 µg/ml of, kanamycin, gentamycin, tetracyclin, IPTG and Bluo-Gal. Individual white colonies were inoculated into 5-ml LB medium with the same antibiotics, cultured overnight (37°C, orbital shaking at 180 rpm), and the bacmid produced was isolated. The baculovirus was produced by transfecting 1 ml of Sf9 insect cells (0.9 × 10^6^ cells/ml in Grace medium, Invitrogen) with 45 µg of recombinant bacmid carrying the DNA-encoding sequences for Wuhan-Hu-1, S:D614G or [S:A222V + S:D614G] variants (proven by PCR), using 0.65 % Fugene (Promega). After 5 hr of plate incubation at 27°C, Grace medium was replaced by Sf900 medium (Invitrogen) containing 0.1% Pluronic F-68, 50 U/ml penicillin and 50 µg/ml streptomycin and cells were cultured for 4-day at 27°C. At the end of the 4 days, the culture medium containing low baculoviral titers, was collected by centrifugation and used to infect a suspension of 1.5 × 10^6^ Sf9 cells/ml by diluting it 60 fold in the cell suspension. After 96-hr culturing (27°C, orbital shaking at 125 rpm) and centrifugation, the high-titre baculovirus supernatant produced was collected and used to infect Sf9 cells at a density of 3 × 10^6^ Sf9 cells/ml by 1:8 dilution of the supernatant virus stock. After 4 days of culture, the produced variant of SARS-CoV-2 spike protein was purified from the medium, where the protein was secreted, by spinning out the cells (1 hour, 19,000 × g) and bringing the pH to neutrality by addition of 10 % volume of a solution contaning 0.5 M Tris-HCl pH 7.4, 60 mM KCl, 2.8 mM NaCl.

Subsequent steps were carried out at 4°C unless indicated. A peristaltic pump was used to apply the culture supernatant (700-1000 mL) to a 5 mL-HisTrap^TM^ Excel column (Cytiva) equilibrated with 20 mM Na-Hepes pH 7.2, 150 mM NaCl (buffer A). After mounting the column to al FPLC, and washing with buffer A supplemented with 25 mM imidazole, SARS-CoV-2 S protein was eluted with 0.5 M imidazole in buffer A, collecting fractions of 2 ml. Fractions containing SARS-CoV-2 S protein (shown by SDS-PAGE) were pooled and concentrated to 0.5 ml by centrifugal ultrafiltration (100-kDa cutoff membrane, Amicon Ultra; Ultracel). The SARS-CoV-2 S protein was then purified by size exclusion chromatography (SEC) using a Superdex 200 Increase 10/300 GL column (Cytiva) pre-equilibrated with 10 mM sodium-Hepes pH 7.2, 150 mM NaCl (SEC buffer), also collecting 0.2-ml fractions, of which those containing SARS-CoV-2 S protein were aliquoted, flash-frozen in liquid nitrogen and stored at -80 °C. SARS-coV-2 S protein concentration was determined spectrophotometrically from the optical absorption at 280 nm, using a sequence-deduced (EXPASY ProtParam tool, https://www.expasy.org) mass extinction coefficient (E 1%) of 10.29 g^-1^L cm^-1^.

ACE2 was produced and purified essentially as described for SARS-CoV-2 protein S, except for the fact that before SEC, the polyhistidine tag was removed by digestion with TEV protease (37.5 µg per each mg of ACE2, overnight, 4°C) within a dialysis bag dialyzed against SEC buffer supplemented with 5 mM EDTA and 0.5 mM mercaptoethanol. Nontagged ACE2 was isolated by an additional Ni-affinity chromatography step, using a 1 mL-HisTrap^TM^ Excel column (Cytiva). In this step, the untagged protein was collected in the initial fraction not retained by the column (application and 2-mL wash). After concentration by centrifugal ultrafiltration (30-kDa cutoff membrane), the protein was further purified by SEC as described for SARS-CoV-2 protein S, quantifying ACE2 by its optical absorption at 280 nm (mass extinction coefficient, 21.9 g^-1^ L cm^-1^ was considered as mass extinction coefficient (E1%). The absence of the polyhistidine tag in the purified protein was verified by western blot using antihistidine antibodies.

### In vitro functional assays

#### Thermal shift assays

Thermofluor assays (Vedadi et al. 2006) were performed in 20 μl of a solution of 0.1 mg/ml SARS-CoV-2 spike protein (Wuhan-Hu-1, S:D614G, or [S:A222V + S:D614G] variants), in 10 mM Na Hepes pH 7.3, 150 mM NaCl and a 1:1,000 dilution of SYPRO Orange (commercial preparation from Invitrogen, Carlsbad, CA), kept in sealed microwell plates. A real-time PCR instrument (7500 model from Applied Biosystems, Thermo Fisher Scientific, Alcobendas, Madrid, Spain) was used to monitored the increase in SYPRO Orange fluorescence (excitation at 488 nm; emission at 610 nm) with temperature increase at a ramp of 1° C /min. The samples were preincubated 20 minutes at 37°C prior to performing the assays (Edwards et al. 2021). Plots, curve fittings and numerical calculations were performed with the program GraphPad Prism 5 (GraphPad Software, San Diego, CA, USA). Results given in Table 1 correspond to the mean and standard error for six replicate assays carried in groups of three replicates in two independent experiments.

#### Biolayer Interferometry assays

Binding assays of ACE2 to His-tagged S proteins were performed in the Octet K2 instrument (ForteBio). The assays were carried out at 28°C, shaking was kept at 1,000 rpm and the solution was 20 mM Hepes, 150 mM NaCl, 50 μM EDTA, 10 mM imidazole and 0.005% Tween 20 (buffer AB). His-tagged Spike proteins at 0.1 μg/ml in solution AB were immobilized (240 s) on Ni-NTA biosensors hydrated in the same buffer, yielding a typical signal of ∼2 nm. After another 240 sec equilibration of the biosensors with buffer AB to get a stable baseline, binding was assayed by adding serial 1:2 dilutions of ACE2 in the range 250-3.9 nM. Both the association and the dissociation phases lasted 120 s each. Negative drift controls were carried out without ACE2 and a 1:1 binding model was used for data fitting.

### Cryo-electron microscopy sample preparation

#### Cryo-EM Data Acquisition

Purified SARS-CoV-2 spike samples in 10mM Hepes pH7.2 and 150mM NaCl (3 μL at 0.62-0.75mg/ml) were kept at 4C until their application onto QUANTIFOIL R 1.2/1.3 Cu:300-mesh grids (QUANTIFOIL). The grids were vitrified using a Leica GP automatic vitrification robot (Leica). Chamber conditions were set at 10°C and 95% relative humidity. Grids were glow discharged for 30 seconds prior to application of the samples. Data were collected on a FEI Talos electron microscope operated at 200 kV and images recorded on a FEI Falcon III detector operating in electron counting mode. A total of 4,841 and 2,492 movies were recorded for [S:A222V + S:D614G] and S:D614G, respectively; at a calibrated magnification of 120,000x, yielding a pixel size of 0.85 Å on the specimen. Each movie comprises 60 frames with an exposure rate of 0.54 e-/Å^2^ per frame, with a total exposure time of 20 s and an accumulated exposure of 32.4 e-/Å^2^. Data acquisition was performed with EPU Automated Data Acquisition Software for Single Particle Analysis (Thermo Fisher Scientific) at -0.3 μm to -3.5 μm defocus.

#### Image Processing

All image processing steps were performed inside Scipion (de la Rosa-Trevín et al. 2016). Movies were motion-corrected and dose weighted with MOTIONCOR2 (Zheng et al. 2017). Aligned, non-dose weighted micrographs were then used to estimate the contrast transfer function (CTF) with GCTF (Zhang 2016) and CTFFIND4.1 (Rohou and Grigorieff 2015). The Scipion CTF consensus protocol (Sorzano et al. 2021) was used to select 4,330 and 1,831 micrographs for, respectively, [S:A222V + S:D614G] and S:D614G (**Supplementary Figure 6** and **7**). Micrographs were then automatically picked using gautomatch and crYOLO (Wagner et al. 2019). Following the application of the Scipion picking consensus protocol 509,466 and 268,763 particles were extracted for [S:A222V + S:D614G] and S:D614G, respectively. From now on pairs of values will be given for each step in the order corresponding to these two variants of S. Images were binned to 1.40 Å/px from this step onwards. 2D classification was performed in cryoSPARC (Punjani et al. 2017) and 310,162 and 165,304 particles were selected. The CryoSPARC initial model protocol was then used to generate and classify the particles into 4 classes, with and without imposing C3 symmetry. Several datasets being more or less restrictive with the number of classes and thus particles selected, were refined using non-uniform refinement in cryoSPARC with no symmetry application, to resolutions of 3.4 Å and 4.2 Å based on the gold-standard (FSC = 0.143) criterion. Several datasets were also selected for applying the C3 symmetry approach, being refined to resolutions of 3.2 Å and 3.7 Å. The dataset with the highest number of particles was then 3D classified in Relion3.1 (Scheres 2012) using the symmetry relaxation protocol (Ilca et al. 2019; Goetschius et al. 2019), which relaxes the dominant symmetry by considering multimodal priors centered at the symmetry related positions. A 30 Å low-pass filtered reconstruction with no symmetry resulting from a randomly selected subset of 10,000 symmetry broken particles (Goetschius et al. 2019) was used as the initial model for the 3D classification. This protocol was repeated three times with the same parameters and to overcome the variability in the results, the Scipion 3D classification consensus protocol was used to better characterize the different conformations (Sorzano et al. 2021). An initial model was generated for each class/conformation and subsequently refined to a resolution of 3.3-3.4 Å ([S:A222V + S:D614G]-1-up), 6.6-6.8 Å ([S:A222V + S:D614G]-2-up), 7.0 Å ([S:A222V + S:D614G]-3-down), 4.1-4.2 Å (S:D614G-1-up) and 8.0-8.8 Å (S:D614G-2-up). The resulting maps were sharpened with DeepEMhancer (Sanchez-Garcia et al. 2021).

#### Model building and refinement

For model building of [S:A222V + S:D614G] in the 1-up conformation ([S:A222V + S:D614G]-1-up), we started with a deposited PDB file for the S protein (PDB: 7BNN). After a simple fitting of 7BNN to the [S:A222V + S:D614G]-1-up map using Chimera (Pettersen et al. 2004), a rigid-body docking of different regions of each monomer for keeping the secondary structure of Spike (14-293; 294-319 plus 592-699; 320-330; 331-529 plus 530-591; 734-775; 944-962; 963-989; 990-1027) was done using Coot (Emsley et al. 2010). Manual building, deletion of disordered loops in the model as well as deletion and/or addition of glycosylations were done in Coot followed by several rounds of refinement in Refmac (Brown et al. 2015). This process was repeated until acceptable refinement metrics were obtained. The refined [S:A222V + S:D614G]-1-up model was used as the starting point for model building of S:D614G- 1-up. Again, a simple fitting with Chimera using our [S:A222V + S:D614G]-1-up model and the S:D614G-1-up map was done, followed by a rigid-body docking of the same regions as described above. After several rounds of refinement in Refmac and model building in Coot, acceptable refinement metrics were obtained. In both models, the lower resolution in a few regions made it difficult to define the structures for some loops (residues 621-641 and 676-689 for S1 and residues 827-854 for S2). Thus, all these procedures generated two models, one for S:D614G-1-up and another one for [S:A222V + S:D614G]-1-up, and both were used for the structural studies performed in this work.

### Molecular dynamics simulations

#### System preparation

The closed, 3-down and open, 1-up states of the glycan-free ectodomain (head-only) of the SARS-CoV-2 Spike glycoprotein were retrieved from the COVID-19 archive (https://www.charmm-gui.org/?doc=archive&lib=covid19) and used to generate the closed and open states for the 20E (EU1) mutant. The two models were built respectively from the cryo-EM structure of the wild type Spike glycoprotein in its 3-down (named “DDD”; PDB: 6VXX) (Walls et al. 2020) and 1-up (named “UDD”; PDB: 6VSB) (Wrapp et al. 2020) conformations. Point mutations at position 222 (A-->V) of the S1-NTD domain (residues 13-305) and 614 (D-->G) of the S1/S2 furin cleavage site were introduced by using the mutagenesis tool of PyMol 2.0 (glycan-free) or an in-house topology editing tool (glycosylated; see gitlab.com/KomBioMol/gromologist). For comparative purposes, the S:D614G mutant was also simulated. Accordingly, four systems named DDDS:D614G, DDD[S:A222V + S:D614G], UDDS:D614G and UDD[S:A222V + S:D614G] and other two models named *gly*-UDDS:D614G and *gly*-UDD[S:A222V + S:D614G] for the glycosylated spike were examined. The same protocol was applied to prepare the cryo-EM (glycan-free) structures, *cryo*-UDDS:D614G and *cryo*-UDD[S:A222V + S:D614G] which were also subjected to MD simulations. The Amber ff99sb-ildn force field (Hornak et al. 2006; Lindorff-Larsen et al. 2010) was used to simulate the protein.

Standard protonation states at pH 7.4 were adopted for ionizable residues. The same N- and O-glycosylation profile, consisting of 22 N-glycans and 1 O-glycan per each monomer as described in the work of Woo H. et al. 2020 (Woo H. et al. 2020), was applied for the simulation. Finally, for all the simulated systems, the protein was enclosed in a truncated octahedron box, solvated with TIP3P (Jorgensen et al. 1983) water molecules and neutralized by adding Na^+^ or K^+^/Cl^-^ counterions (Joung and Cheatham 2008). A detailed description of all the MD systems studied in this work is reported in **Supplementary Table 2**.

#### MD simulations

Classical MD simulations were carried out with Gromacs v. 2020 (Abraham et al. 2015). The systems were energy minimized by applying 50,000 steps of the steepest descent algorithm followed by 5,000 steps of conjugate gradient algorithm. 1 ns of MD simulation in the NVT ensemble (Bussi et al. 2007) was run using the velocity-rescaling thermostat and a time coupling constant of 0.1 ps to heat the systems to the production temperature of 300 K. Positional restraints with a force constant of 1,000 kJ mol^−1^ mn^−2^ were applied to the protein backbone atoms to avoid unnatural distortions during heating. Equilibration to the target temperature was then accomplished in two steps consisting of (i) 5 ns of unrestrained equilibration in the NPT ensemble using Berendsen thermostat (0.1 ps time coupling constant) and barostat (0.5 ps time coupling constant) and (ii) 5 ns of unrestrained MD simulation in the NPT ensemble by using Berendsen thermostat (0.1 ps time coupling constant) and Parrinello-Rahman barostat (0.5 ps time coupling constant) (Parrinello 1981). For the glycan-free systems, 300 ns of NPT MD production were run for the 3-down (DDD) state in periodic boundary conditions by using the Parrinello–Rahman barostat. For the 1-up (UDD) state, a longer MD run (500 ns) was performed to enhance conformational sampling. All the simulations were run in triplicate, leading to a global simulation time of 2.4 μs (0.9 + 1.5 μs; **Supplementary Table 2**) for the S:D614G and [S:A222V + S:D614G] mutants. For the fully-glycosylated systems based on 6VSB and the cryo-EM (glycan-free) systems, respectively, 400 and 200 ns of NPT MD production were run for the UDD models of both the S:D614G and [S:A222V + S:D614G] mutants, leading to a global simulation time of 1.2 μs (0.8 + 0.4 μs; **Supplementary Table 2**). In all the cases, the LINCS method (Hess et al. 1998) was applied to constraint bonds involving hydrogen atoms. A cut-off of 1.2 nm was used to treat short-range nonbonded interactions, whereas the PME method was applied to manage long-range electrostatic interactions (Darden et al. 1993). A time step of 2 fs was applied to collect trajectories during the simulation.

#### Convergence assessment

Root-mean-square deviation (RMSD) analysis was carried out to evaluate the structural stability and convergence of the simulated systems. The *gmx rms* tool of gromacs 2020 was used to calculate the 1D positional rmsd for the protein Cα atoms over time. The RMSD profiles (**Supplementary Figure 13**) supported the structural stability for each of the three replicas for the S:D614G and [S:A222V + S:D614G] mutants in the closed conformation (**Supplementary Figure 13a-b**), as noted in average RMSD values ranging between 0.2 and 0.4 nm with respect to the starting, energy minimized structure. Slightly higher deviations (up to 0.7 nm) were observed for the open state (**Supplementary Figure 13c-d**) although all the simulated systems globally reached structural convergence after 150 ns of MD simulation. The same analysis was done on the MD trajectories for the mutants (i) based on 6VSB and on (ii) the cryo-EM (glycan-free) structures and reported in **Supplementary Figure 13e-f**.

#### Principal components analysis (PCA)

The internal collective motions of the principal S1 subunit for the S:D614G and [S:A222V + S:D614G] mutants of the Spike protein were analysed by calculating the positional covariance matrix for the Cα backbone atoms. For this analysis, all trajectories were re-aligned on the protein Cα atoms of the central helixes of the trimeric S2 domains (Cα atoms for residues 718A-994A, 718B-994B and 718C-994C). For each simulated glycan-free system (S:D614G and [S:A222V + S:D614G]), all frames across all replicas for the concatenated DDD and UDD trajectories were considered. The gromacs tool *gmx covar* was used to extract the eigenvalues and eigenvectors from the 2.4 and 0.4 μs-long MD trajectories, and *gmx anaeig* was used to analyse and plot the eigenvectors. For comparative purposes, the [S:A222V + S:D614G] mutant was always projected on the eigenvectors of the reference S:D614G mutant. For consistency, MD trajectories collected from cryo-EM and glycosylated mutants were also aligned on the previously cited ensemble. Another PCA focused on residues pertaining to the three RBD (residues 330A-530A, 330B-530B, 330C-530C) was also performed and used to calculate continuous density distributions along the first eigenvector (PC1^RBD^). Results from this analysis were compared with those obtained for projection along the first eigenvector (PC1^S1^) for the whole S1 (residues 30A-650A, 30B-650B, 30C-650C).

#### Dynamic network analysis

The dynamic network analysis was performed using an implementation by Melo *et al*. (Melo et al. 2020), following a standard set of parameters, i.e. a contact cut-off of 4.5 Å and contact persistence of 0.5. Since each system consisted of the same selection (open RBD + the neighboring NTD), the reported values of betweenness centrality are absolute rather than normalized within a system. ParmEd (Shirts et al. 2017) was used to convert between trajectory and topology formats as required for the analysis.

#### Mutational free energy analysis

Free energy calculations were performed using the non-equilibrium alchemical protocol based on the Crooks theorem (Crooks 1998). The PMX library was used to introduce the modifications (Gapsys et al. 2015), and an in-house script (gitlab.com/KomBioMol/crooks) was used to run and analyse the free energy calculations. Bennett acceptance ratio (BAR) was used to determine the free energy values (Bennett 1976; Shirts et al. 2003). Starting points for individual non-equilibrium runs were sampled from 200-ns equilibrium simulations of the respective Spike mutant in the 1-up conformation (prepared as described above), and calculations were repeated for each of the three chains to compare the effect of mutations on the thermodynamic preference for the open, 1-up state.

### PCA of existing S:D614G structures

24 structures from six recent structural studies of the S:D614G spike were analysed together with our two new cryo-EM structures (**Supplementary Table 4**). More structural models were available in the PDB but were not included as they came from earlier rounds of classification as structures that were included (PDB: 7KDK and 7KDL) (Gobeil et al. 2021) or had a poorly resolved S1 subunit (PDB: 6XS6) (Yurkovetskiy et al. 2020). Principal component analysis was performed using the latest version of the Protein Dynamics (*ProDy*) application programming interface (API) (Zhang et al. 2021) from GitHub as follows. Only Cα atoms were included for efficiency. A structural ensemble was created using a WT 1-up structure (PDB: 6VSB) as a reference for initial alignment, where the chains were relabelled to match our cryo-EM structures. Residues were matched with the default method and chains with a custom matching function using the chain order correspondences in the second last column of **Supplementary Table 4**, based on a visual analysis in PyMOL. Initial alignment and superposition to the reference was then followed by iterative superposition until the mean structure converged. According to the protocol used for the PCA of our MD simulations, structures were then iteratively superposed again using only the Cα atoms of core helices of the S2 trimer. The ensemble was then trimmed to exclude residues only found in some structures. Positional covariance matrices were calculated based on the single subunit related to RBD opening as done for the MD simulations as well as the whole spike. In the case of the whole spike, covariance matrices were calculated using both the whole ensemble and a subset lacking the three 2-up structures. The three resulting covariance matrices were each decomposed to yield the first 20 eigenvectors describing the most significant modes of structural variation with eigenvalues corresponding to the extent of variance covered. One to three modes, which contributed >90 % of the variance, were considered the principal components (PCs). Square fluctuations were calculated from all modes as eigenvalue-weighted dot products of corresponding eigenvectors. Matrices of directional overlaps were calculated as correlation cosines. For comparison with the single subunit PCA, after slicing the eigenvectors from the whole spike were sliced to only include elements corresponding to the single subunit. Conformations along individual eigenvectors were generated using the Normal Mode Wizard (NMWiz) (Bakan and Bahar 2011) in VMD (Humphrey et al. 1996) and 40 conformers with an RMSD up to 4 Å in both directions were used for making figures.

### Data Availability

All the analyzed 3,227,996 SARS-CoV-2 genomes (collected from May 2020 to October 2021) were retrieved from the GISAID platform (https://www.gisaid.org).

The atomic coordinates for [S:A222V + S:D614G]-1-up and S:D614G-1-up were deposited in the Protein Data Bank with codes 7QDG and 7QDH, respectively. The [S:A222V + S:D614G]-1-up, [S:A222V + S:D614G]-2-up, [S:A222V + S:D614G]-3-down, S:D614G-1-up and S:D614G-2-up cryo-EM density maps were deposited in the EM Data Bank with codes EMD-13916, EMD-13917, EMD-13918, EMD-13919 and EMD-13920, respectively. The motion corrected micrographs for [S:A222V + S:D614G] are being deposited in EMPIAR.

Aligned MD trajectories for all the glycan-free and glycosylated S:D614G and [S:A222V + S:D614G] mutants simulated in this study are available at: bioexcel-cv19-dev.bsc.es/#/browse?search=Spike%20mutation%20A222V

## Acknowledgements

We want to particularly acknowledge the patients and the Consorcio Hospital General de Valencia Biobank integrated in the Valencian Biobanking Network for their collaboration in providing the convalescent serum samples. We acknowledge access to the cryoEM CNB-CSIC facility in the context of the CRIOMECORR project (ESFRI-2019-01-CSIC-16), and in particular the help of its staff. CPU time was partially provided by the PL-Grid Infrastructure.

This research work was supported by the European Commission–NextGenerationEU through the CSIC Global Health Platform. Additionally, authors would like to acknowledge economic support from the Spanish Ministry of Science and Innovation through Grants: PID2019-104757RB-I00 funded by MCIN/AEI/ 10.13039/501100011033, RTI2018-094399-A-I00, and “ERDF A way of making Europe”, by the “European Union”, Grant SEV 2017-0712 funded by MCIN/AEI /10.13039/501100011033, the “Comunidad Autónoma de Madrid” through Grant: S2017/BMD-3817, and the European Union (EU) and Horizon 2020 through grants: Marie-Curie Fellowship EnLaCES (MSCA IF 2020, Proposal: 101024130) (to **JK**), HighResCells (ERC - 2018 - SyG, Proposal: 810057), and iNEXT-Discovery (Proposal: 871037). **AM**, **VR**, **JB** and **JLL** are funded by CIBERER-ISCIII (proposal: COV20/00437), Fondo Supera COVID-19 (proposal: CSIC- COVID19-082), Banco Santander (Proposal: BlockAce), and CSIC PTI Salud Global (Proposal: 202080E110). **VR** is funded by the Spanish Ministry of Science and Innovation through Grant PID2020-120322RB-C21. **IC** is funded by project PID2019-104477RB-100, Fondo COVID COV20/00140 and ERC CoG 101001038. **MC** is funded by the RyC program from the Spanish Ministry of Science and Innovation, the Generalitat Valenciana (SEJI/2019/011).

## Author Contributions

These authors contributed equally: Tiziana Ginex (**TG**), Clara Marco-Marín (**CMM**), Miłosz Wieczór (**MW**), Carlos P. Mata (**CPM**), James Krieger (**JK**). **TG** coordinated and performed the MD simulations, analysed the results and contributed to write the manuscript. **CMM** helped to make the samples for *in vitro* functional assays and preparation of cryo-EM grids, to build the structural models, to analyse the data, to supervise the work and to write the manuscript. **MW** performed the MD simulations and the free energy calculations, analysed the results and wrote the manuscript. **CPM** performed the cryo-EM image processing, analysed the results and contributed to write the manuscript. **JK** performed the PCA of experimental S:D614G structures, and helped with analysing the MD simulations and writing the manuscript. **MLLR** prepared the cryo-EM grids, helped to collect the cryo-EM data, to build the structural models, to analyse the data, and to write the manuscript. **CFG** produced the pseudotype virus, performed the pseudotype neutralization assays, and analyzed the results. **PRR** prepared the global sequence set, performed sequence and phylogenetic analysis, and wrote the manuscript. **RM** collected the cryo-EM data and was involved in the cryo-EM image processing. **COSS** and **MM** were involved in the cryo-EM image processing. **NG** helped to make the samples for in vitro functional assays and formation of cryo-EM grids. **AFN** performed the thermal shift assay. **SZC**, **RGR** and **CSF** performed the biolayer interferometry assays. **JB**, **VR**, and **AM** supervised the work and helped to write the manuscript. The **IBV-Covid19-Pipeline** produced the ACE2 protein used in biolayer interferometry assays. **RG** supervised the neutralization assay and helped to write the manuscript.

**IC** supervised the work and helped to write the manuscript. **CG** supervised the work and helped in the preparation of the manuscript. **MC** supervised sequences and phylogenetic analysis and wrote the manuscript. **MO** supervised the work and helped to write the manuscript. **JLL** supervised the work and helped to build the structural models, to analyse the data and to write the manuscript. **JMC** coordinated the work among the laboratories, the analysis of the data and the writing of the manuscript.

All authors read and approved the final manuscript.

## Consortia

### The IBV-Covid19-Pipeline

Laura Villamayor (orcid.org/0000-0002-7654-3306), Carolina Espinosa, Anmol Adhav (orcid.org/0000-0002-3504-8675), Maria del Pilar Hernández-Sierra, Rafael Ruiz-Partida (orcid.org/0000-0002-1696-6753).

### Corresponding author

Correspondence to José-Maria Carazo, Centro Nacional de Biotecnología (CNB-CSIC), 28049 Madrid, Spain; ORCID: orcid.org/0000-0003-0788-8447.

## Supplementary Information

Supplementary Figures 1-13, Movie 1 and Supplementary Tables 1-5.

## References

1. Abraham, M. J., T. Murtola, R. Schulz, S. Pall, J. C. Smith, B. Hess, and E. Lindahl. 2015. “GROMACS: High performance molecular simulations through multi-level parallelism from laptops to supercomputers.” SoftwareX1-2:19–25. DOI: 10.1016/j.softx.2015.06.001.

2. Amaro, R. E., and A. J. Mulholland. 2020. “A Community Letter Regarding Sharing Biomolecular Simulation Data for COVID-19.” J Chem Inf Model. 60 (6): 2653–2656. DOI: 10.1021/acs.jcim.0c00319.

3. Bakan, A., and I. Bahar. 2011. “Computational generation inhibitor-bound conformers of p38 MAP kinase and comparison with experiments.” Pac Symp Biocomput., 181–92. DOI: 10.1142/9789814335058_0020.

4. Bennett, C. H. 1976. “Efficient estimation of free energy differences from Monte Carlo data.” J Comput Phys. 22 (2): 245–268. DOI: 10.1016/0021-9991(76)90078-4.

5. Benton, D. J., A. G. Wrobel, C. Roustan, A. Borg, P. Xu, S. R. Martin, P. B. Rosenthal, J. J. Skehel, and S. J. Gamblin. 2021. “The effect of the D614G substitution on the structure of the spike glycoprotein of SARS-CoV-2.” Proc Natl Acad Sci U S A. 118 (9): e2022586118. DOI: 10.1073/pnas.2022586118.

6. Brown, A., F. Long, R. A. Nicholls, J. Toots, P. Emsley and G. Murshudov. 2015. “Tools for macromolecular model building and refinement into electron cryo-microscopy reconstructions”. Acta Crystallogr D Biol Crystallogr. 71(Pt 1): 136–153. DOI: 10.1107/S1399004714021683.

7. Bussi, G., D. Donadio, and M. Parrinello. 2007. “Canonical sampling through velocity rescaling.” J Chem Phys. 126 (1): 014101. DOI: 10.1063/1.2408420.

8. Casalino, L., Z. Gaieb, J. A. Goldsmith, C. K. Hjorth, A. C. Dommer, A. M. Harbison, C. A. Fogarty, et al. 2020. “Beyond Shielding: The Roles of Glycans in the SARS-CoV-2 Spike Protein.” ACS Cent Sci. 6 (10): 1722–1734. DOI: 10.1021/acscentsci.0c01056.

9. Chen, M., and S. J. Ludtke. 2021. “Deep learning-based mixed-dimensional Gaussian mixture model for characterizing variability in cryo-EM.” Nat Methods. 18:930–936. DOI:10.1038/s41592-021-01220-5.

10. Crooks, G. E. 1998. “Nonequilibrium Measurements of Free Energy Differences for Microscopically Reversible Markovian Systems.” J. Stat. Phys. 90:1481–1487. DOI:10.1023/A:1023208217925.

11. Darden, T., D. York, and Lee Pedersen. 1993. “Particle mesh Ewald: An N⋅log(N) method for Ewald sums in large systems.” J Chem Phys. 98:10089. DOI:10.1063/1.464397.

12. de la Rosa-Trevín, J. M., A. Quintana, L. Del Cano, A. Zaldívar, I. Foche, J. Gutiérrez, J. Gómez-Blanco, et al. 2016. “Scipion: A software framework toward integration, reproducibility and validation in 3D electron microscopy.” J Struct Biol. 195 (1): 93–9. DOI: 10.1016/j.jsb.2016.04.010.

13. Elbe, S., and G. Buckland-Merrett. 2017. “Data, disease and diplomacy: GISAID’s innovative contribution to global health.” Glob Chall. 1:33–46. DOI: 10.1002/gch2.1018.

14. Emsley, P., B. Lohkamp, W. G. Scott, and K. Cowtan. 2010. “Features and development of Coot.” Acta Crystallogr D Biol Crystallogr. 66 (Pt 4): 486–501. DOI: 10.1107/S0907444910007493.

15. Gapsys, V., S. Michielssens, D. Seeliger, and B. L. de Groot. 2015. “pmx: Automated protein structure and topology generation for alchemical perturbations.” J Comput Chem. 36 (5): 348–54. DOI: 10.1002/jcc.23804.

16. Gobeil, S. M., K. Janowska, S. McDowell, K. Mansouri, R. Parks, K. Manne, V. Stalls, et al. 2021. “D614G Mutation Alters SARS-CoV-2 Spike Conformation and Enhances Protease Cleavage at the S1/S2 Junction.” Cell Rep. 34 (2): 108630. DOI: 10.1016/j.celrep.2020.108630.

17. Goetschius, D. J., H. Lee, and S. Hafenstein. 2019. “CryoEM reconstruction approaches to resolve asymmetric features.” Adv Virus Res. 105:73–91. DOI: 10.1016/bs.aivir.2019.07.007.

18. Hess, B., H. Bekker, J. C. Herman, H. J. Berendsen, and J. G. Fraaije. 1998. “LINCS: A Linear Constraint Solver for molecular simulations.” J Comput Chem. 18:1463–1472. DOI: 10.1002/(SICI)1096-987X(199709)18:123.0.CO;2-H.

19. Hodcroft, E. B., M. Zuber, S. Nadeau, T. G. Vaughan, K. H. Crawford, C. L. Althaus, M. L. Reichmuth, et al. 2021. “Spread of a SARS-CoV-2 variant through Europe in the summer of 2020.” Nature. 595 (7869): 707–712. DOI: 10.1038/s41586-021-03677-y.

20. Hornak, V., R. Abel, A. Okur, B. Strockbine, A. Roitberg, and C. Simmerling. 2006. “Comparison of multiple Amber force fields and development of improved protein backbone parameters.” Proteins. 65 (3): 712–25. DOI: 10.1002/prot.21123.

21. Hsieh, C. L., J. A. Goldsmith, J. M. Schaub, A. M. DiVenere, H. C. Kuo, K. Javanmardi, K. C. Le, et al. 2020. “Structure-based design of prefusion-stabilized SARS-CoV-2 spikes.” Science. 369 (6510): 1501–1505. DOI: 10.1126/science.abd0826.

22. Humphrey, W., A. Dalke, and K. Schulten. 1996. “VMD: visual molecular dynamics.” J Mol Graph. 14 (1): 33–8, 27–8. DOI: 10.1016/0263-7855(96)00018-5.

23. Ilca, S. L., X. Sun, K. El Omari, A. Kotecha, F. de Haas, F. DiMaio, J. M. Grimes, D. I. Stuart, M. M. Poranen, and J. T. Huiskonen. 2019. “Multiple liquid crystalline geometries of highly compacted nucleic acid in a dsRNA virus.” Nature. 570 (7760): 252–256. DOI: 10.1038/s41586-019-1229-9.

24. Jorgensen, W. L., J. Chandrasekhar, and J. D. Madura. 1983. “Comparison of simple potential functions for simulating liquid water.” J Chem Phys. 79:926. DOI: 10.1063/1.445869.

25. Joung, I. S., and T. E. Cheatham. 2008. “Determination of alkali and halide monovalent ion parameters for use in explicitly solvated biomolecular simulations.” J Phys Chem B. 112 (30): 9020–41. DOI: 10.1021/jp8001614.

26. Korber, B., W. M. Fischer, S. Gnanakaran, H. Yoon, J. Theiler, W. Abfalterer, N. Hengartner, et al. 2020. “Tracking Changes in SARS-CoV-2 Spike: Evidence that D614G Increases Infectivity of the COVID-19 Virus.” Cell. 182 (4): 812–827.e19. DOI: 10.1016/j.cell.2020.06.043.

27. Lan, J., J. Ge, J. Yu, S. Shan, H. Zhou, S. Fan, Q. Zhang, et al. 2020. “Structure of the SARS-CoV-2 spike receptor-binding domain bound to the ACE2 receptor.” Nature. 581 (7807): 215–220. DOI: 10.1038/s41586-020-2180-5.

28. Lindorff-Larsen, K., S. Piana, K. Palmo, P. Maragakis, J. L. Klepeis, R. O. Dror, and D. E. Shaw. 2010. “Improved side-chain torsion potentials for the Amber ff99SB protein force field.” Proteins. 78 (8): 1950–8. DOI: 10.1002/prot.22711.

29. Mashayekhi, G., J. Vant, A. Singharoy, and A. Ourmazd. 2021. “Energy Landscape of the SARS-CoV-2 Reveals Extensive Conformational Heterogeneity.” BioRxiv [Preprint]. DOI:10.1101/2021.05.11.443708.

30. Melero, R., C. O. Sorzano, B. Foster, J. L. Vilas, M. Martínez, R. Marabini, E. Ramírez-Aportela, et al. 2020. “Continuous flexibility analysis of SARS-CoV-2 spike prefusion structures.” IUCrJ. 7 (Pt 6): 1059–69. DOI: 10.1107/S2052252520012725.

31. Melo, M. C., R. C. Bernardi, C. de la Fuente-Nunez, and Z. Luthey-Schulten. 2020. “Generalized correlation-based dynamical network analysis: a new high-performance approach for identifying allosteric communications in molecular dynamics trajectories.” J Chem Phys. 153 (13): 134104. DOI: 10.1063/5.0018980.

32. Ozono, S., Y. Zhang, H. Ode, K. Sano, T. S. Tan, K. Imai, K. Miyoshi, et al. 2021. “SARS-CoV-2 D614G spike mutation increases entry efficiency with enhanced ACE2-binding affinity.” Nat Commun. 8 (12): 848. DOI: 10.1038/s41467-021-21118-2.

33. Page, A. J., B. Taylor, A. J. Delaney, J. Soares, T. Seemann, J. A. Keane, and S. R. Harris. 2016. “SNP-sites: rapid efficient extraction of SNPs from multi-FASTA alignments.” Microb Genom. 2:e000056. DOI:10.1099/mgen.0.000056.

34. Pang, Y. T., A. Acharya, D. L. Lynch, A. Pavlova, and J. C. Gumbart. 2021. “SARS-CoV-2 spike opening dynamics and energetics reveal the individual roles of glycans and their collective impact.” BioRxiv [Preprint]. DOI: 10.1101/2021.08.12.456168.

35. Parrinello, M. 1981. “Polymorphic transitions in single crystals: A new molecular dynamics method.” J Appl Phys.52:7182. DOI:10.1063/1.328693.

36. Pettersen, E. F., T. D. Goddard, C. C. Huang, G. S. Couch, D. M. Greenblatt, E. C. Meng, and T. E. Ferrin. 2004. “UCSF Chimera--a visualization system for exploratory research and analysis.” J Comput Chem. 25 (13): 1605–12. DOI: 10.1002/jcc.20084.

37. Plante, J. A., Y. Liu, J. Liu, H. Xia, B. A. Johnson, K. G. Lokugamage, X. Zhang, et al. 2021. “Spike mutation D614G alters SARS-CoV-2 fitness.” Nature. 592 (7852): 116–121. DOI: 10.1038/s41586-020-2895-3.

38. Punjani, A., and D. J. Fleet. 2021a. “3D variability analysis: Resolving continuous flexibility and discrete heterogeneity from single particle cryo-EM.” J Struct Biol. 213 (2): 107702. DOI: 10.1016/j.jsb.2021.107702.

39. Punjani, A., and D. J. Fleet. 2021b. “3D flexible refinement: Structure and motion of flexible proteins from cryo-EM.” BioRxiv [Preprint]. DOI: 10.1101/2021.04.22.440893.

40. Punjani, A., J. L. Rubinstein, D. J. Fleet, and M. A. Brubaker. 2017. “cryoSPARC: algorithms for rapid unsupervised cryo-EM structure determination.” Nat Methods. 14 (3): 290–296. DOI: 10.1038/nmeth.4169.

41. Rahnama, S., Irani M. Azimzadeh, M. Amininasab, and M. R. Ejtehadi. 2021. “S494 O-glycosylation site on the SARS-CoV-2 RBD affects the virus affinity to ACE2 and its infectivity; a molecular dynamics study.” Sci Rep. 11 (1): 15162. DOI: 10.1038/s41598-021-94602-w.

42. Ramírez-Aportela, E., Maluenda, D., Fonseca, Y. C., Conesa, P., Marabini, R., Heymann, J. B., Carazo, J. M., and Sorzano, COS. 2021. “FSC-Q: a CryoEM map-to-atomic model quality validation based on the local Fourier shell correlation.” Nat Commun. 12 (1): 42. DOI: 10.1038/s41467-020-20295-w

43. Rohou, A., and N. Grigorieff. 2015. “CTFFIND4: Fast and accurate defocus estimation from electron micrographs.” J Struct Biol. 192 (2): 216–21. DOI: 10.1016/j.jsb.2015.08.008.

44. Ruiz-Rodriguez, P., Francés-Gómez, C., Chiner-Oms, Á., López, M. G., Jiménez-Serrano, S., Cancino-Muñoz, I., Ruiz-Hueso, et al. 2021. “Evolutionary and Phenotypic Characterization of Two Spike Mutations in European Lineage 20E of SARS-CoV-2.” *mBio*, e0231521. DOI: 10.1128/mBio.02315-21.

45. Sanchez-Garcia, R., J. Gomez-Blanco, A. Cuervo, J. M. Carazo, C. O. Sorzano, and J. Vargas. 2021. “DeepEMhancer: a deep learning solution for cryo-EM volume post-processing.” Commun Biol. 4 (1): 874. DOI: 10.1038/s42003-021-02399-1.

46. Scheres, S. H. 2012. “A Bayesian view on cryo-EM structure determination.” J Mol Biol. 415 (2): 406–18. DOI: 10.1016/j.jmb.2011.11.010.

47. Seitz, E., F. Acosta-Reyes, S. Maji, P. Schwander, and J. Frank. 2021. “Geometric machine learning informed by ground truth: Recovery of conformational continuum from single-particle cryo-EM data of biomolecules.”BioRxiv [Preprint]. DOI:10.1101/2021.06.18.449029.

48. Shen, W., S. Le, Y. Li, and F. Hu. 2016. “SeqKit: A Cross-Platform and Ultrafast Toolkit for FASTA/Q File Manipulation.” PLoS One. 11 (10): e0163962. DOI: 10.1371/journal.pone.0163962.

49. Shirts, M. R., E. Bair, G. Hooker, and V. S. Pande. 2003. “Equilibrium free energies from nonequilibrium measurements using maximum-likelihood methods.” Phys Rev Lett. 91 (14): 140601. DOI: 10.1103/PhysRevLett.91.140601.

50. Shirts, M. R., C. Klein, J. M. Swails, J. Yin, M. K. Gilson, D. L. Mobley, D. A. Case, and E. D. Zhong. 2017. “Lessons learned from comparing molecular dynamics engines on the SAMPL5 dataset.” J Comput Aided Mol Des. 31 (1): 147–161. DOI: 10.1007/s10822-016-9977-1.

51. Sorzano, C. O., A. Jiménez-Moreno, D. Maluenda, E. Ramírez-Aportela, M. Martínez, A. Cuervo, R. Melero, et al. 2021. “Image Processing in Cryo-Electron Microscopy of Single Particles: The Power of Combining Methods.” Methods Mol Biol. 2305:257–289. DOI: 10.1007/978-1-0716-1406-8_13.

52. Sorzano, C. O. S., and J. M. Carazo. 2021. “Principal component analysis is limited to low-resolution analysis in cryoEM.” Acta Crystallogr D Struct Biol. 77 (Pt 6): 835–839. DOI: 10.1107/S2059798321002291.

53. Sztain, T., S. H. Ahn, A. Bogetti, L. Casalino, J. A. Goldsmith, E. Seitz, R. S. McCool, et al. 2021. “A glycan gate controls opening of the SARS-CoV-2 spike protein.” Nat Chem. 13 (10): 963–968. DOI: 10.1038/s41557-021-00758-3.

54. van Dorp, L., D. Richard, C. C. Tan, L. P. Shaw, M. Acman, and F. Balloux. 2020. “No evidence for increased transmissibility from recurrent mutations in SARS-CoV-2.” Nat Commun. 11 (1): 5986. DOI: 10.1038/s41467-020-19818-2.

55. Vedadi, M., F. H. Niesen, A. Allali-Hassani, O. Y. Fedorov, P. J. Finerty, G. A. Wasney, R. Yeung, et al. 2006. “Chemical screening methods to identify ligands that promote protein stability, protein crystallization, and structure determination.” Proc Natl Acad Sci U S A. 103 (43): 15835–40. DOI: 10.1073/pnas.0605224103.

56. Wagner, T., F. Merino, M. Stabrin, T. Moriya, C. Antoni, A. Apelbaum, P. Hagel, et al. 2019. “SPHIRE-crYOLO is a fast and accurate fully automated particle picker for cryo-EM.” Commun Biol. 2:218. DOI: 10.1038/s42003-019-0437-z.

57. Walls, A. C., Y. J. Park, M. A. Tortorici, A. Wall, A. T. McGuire, and D. Veesler. 2020. “Structure, Function, and Antigenicity of the SARS-CoV-2 Spike Glycoprotein.” Cell. 181 (2): 281–292.e6. DOI: 10.1016/j.cell.2020.02.058.

58. Watanabe, Y., Z. T. Berndsen, J. Raghwani, G. E. Seabright, J. D. Allen, O. G. Pybus, J. S. McLellan, et al. 2020. “Vulnerabilities in coronavirus glycan shields despite extensive glycosylation.” Nat Commun. 11 (1): 2688. DOI: 10.1038/s41467-020-16567-0.

59. Woo, H., S. J. Park, Y. K. Choi, T. Park, M. Tanveer, Y. Cao, N. R. Kern, et al. 2020. “Developing a Fully Glycosylated Full-Length SARS-CoV-2 Spike Protein Model in a Viral Membrane.” J Phys Chem B. 124 (33): 7128–7137. DOI: 10.1021/acs.jpcb.0c04553.

60. Wrapp, D., N. Wang, K. S. Corbett, J. A. Goldsmith, C. L. Hsieh, O. Abiona, B. S. Graham, and J. S. McLellan. 2020. “Cryo-EM structure of the 2019-nCoV spike in the prefusion conformation.” Science. 367 (6483): 1260–1263. DOI: 10.1126/science.abb2507.

61. Xu, C., Wang, Y., Liu, C., Zhang, C., Han, W., Hong, X., Wang, Y., Hong, Q., Wang, S., Zhao, Q., Wang, Y., Yang, Y., Chen, K., Zheng, W., Kong, L., Wang, F., Zuo, Q., Huang, Z., and Cong, Y. 2021. “Conformational dynamics of SARS-CoV-2 trimeric spike glycoprotein in complex with receptor ACE2 revealed by cryo-EM.” Science advances, 7 (1): eabe5575. DOI: 10.1126/sciadv.abe5575

62. Yang, T. J., P. Y. Yu, Y. C. Chang, and S. D. Hsu. 2021. “D614G mutation in the SARS-CoV-2 spike protein enhances viral fitness by desensitizing it to temperature-dependent denaturation.” J Biol Chem. 297 (4): 101238. DOI: 10.1016/j.jbc.2021.101238.

63. Yurkovetskiy, L., X. Wang, K. E. Pascal, C. Tomkins-Tinch, T. P. Nyalile, Y. Wang, A. Baum, et al. 2020. “Structural and Functional Analysis of the D614G SARS-CoV-2 Spike Protein Variant.” Cell. 183 (3): 739–751.e8. DOI: 10.1016/j.cell.2020.09.032.

64. Zhang, J., Y. Cai, T. Xiao, J. Lu, H. Peng, S. M. Sterling, R. M. Walsh, et al. 2021. “Structural impact on SARS-CoV-2 spike protein by D614G substitution.” Science. 372 (6541): 525–530. DOI: 10.1126/science.abf2303.

65. Zhang, K. 2016. “Real-time CTF determination and correction.” J Struct Biol. 193 (1): 1–12. DOI: 10.1016/j.jsb.2015.11.003.

66. Zhang, S., J. M. Krieger, Y. Zhang, C. Kaya, B. Kaynak, K. Mikulska-Ruminska, P. Doruker, H. Li, and I. Bahar. 2021. “ProDy 2.0: Increased Scale and Scope after 10 Years of Protein Dynamics Modelling with Python.” Bioinformatics. btab187. DOI: 10.1093/bioinformatics/btab187.

67. Zheng, S. Q., E. Palovcak, J. P. Armache, K. A. Verba, Y. Cheng, and D. A. Agard. 2017. “MotionCor2: anisotropic correction of beam-induced motion for improved cryo-electron microscopy.” Nat Methods. 14 (4): 331–332. DOI: 10.1038/nmeth.4193.

68. Zhong, E. D., T. Bepler, B. Berger, and J. H. Davis. 2021. “CryoDRGN: reconstruction of heterogeneous cryo-EM structures using neural networks.” Nat Methods. 18 (2): 176–185. DOI: 10.1038/s41592-020-01049-4.

