## Supplementary Information for "The structural role of SARS-CoV-2 genetic background in the emergence and success of spike mutations: the case of the spike A222V mutation"

**The IBV-Covid19-Pipeline:** Laura Villamayor, Carolina Espinosa, Anmol Adhav, Maria del Pilar Hernández-Sierra, Rafael Ruiz-Partida.

**List of the IBV-Covid19-Pipeline members that did not directly contribute to the manuscript**: Susana Masiá, Francisca Gallego, Monica Escamilla-Aguilar, Antonio Rubio-Del-Campo, Lidia Orea-Ordóñez, Alonso Felipe, Borja Saez-De la Fuente, Guilherme Dim, Alba Iglesias-Ceacero, Francisco Del Caño-Ochoa, Javier Mancheño, Jesus Rodríguez-Díaz, Santiago Ramón-Maiques.

**Table of Content:**

**Supplementary Figures**

*Page S3*	**Supplementary Figure 1.** Temporal distribution of Delta sub-lineages.

*Page S3*	**Supplementary Figure 2.** Protein purification and quality analysis.

*Page S4*	**Supplementary Figure 3.** Thermofluor assays (means of three replicas) of the indicated SARS-CoV-2 S protein mutants.

*Page S5*	**Supplementary Figure 4.** Kinetic analysis of spike interaction to ACE2 measured by biolayer interferometry (BLI).

*Page S6*	**Supplementary Figure 5.** Cryo-EM data of [S:A222V + S:D614G].

*Page S7-8*	**Supplementary Figure 6.** Cryo-EM image processing workflow for S:D614G.

*Page S9-10*	**Supplementary Figure 7.** Cryo-EM image processing workflow for [S:A222V + S:D614G].

*Page S11*	**Supplementary Figure 8.** Cryo-EM of S:D614G.

*Page S12*	**Supplementary Figure 9.** Comparison of the two structures at the region where S:A222V mutation is located.

*Page S13*	**Supplementary Figure 10.** Continuous population densities along the PC1.

*Page S14*	**Supplementary Figure 11.** PCA of whole experimental S:D614G spike structures excluding 2-up conformations.

*Page S15*	**Supplementary Figure 12.** PCA results for whole S:D614G PDB structures including 2-up conformations.

*Page S16*	**Supplementary Figure 13.** RMSD analysis.

**Supplementary Video**

*Page S17*	**Movie 1.** Structural changes from S:D614G to [S:A222V + S:D614G], starting from cryo-EM maps, in two different orientations.

**Supplementary Tables**

*Page S18*	**Supplementary Table 1.** Interaction between RBDs and NTDs domains from different subunits.

*Page S19*	**Supplementary Table 2.** Detailed description of the 3-down (DDD) and 1-up (UDD) models of the SARS-CoV-2 spike mutants simulated in this study.

*Page S20*	**Supplementary Table 3.** Mutational free energy analysis.

*Page S20-21*	**Supplementary Table 4.** Experimental structures of the SARS-CoV-2, S:D614G spike used in PCA analysis.

*Page S21*	**Supplementary Table 5.** Oligonucleotides used in this study.

**References**

**Supplementary Figures**

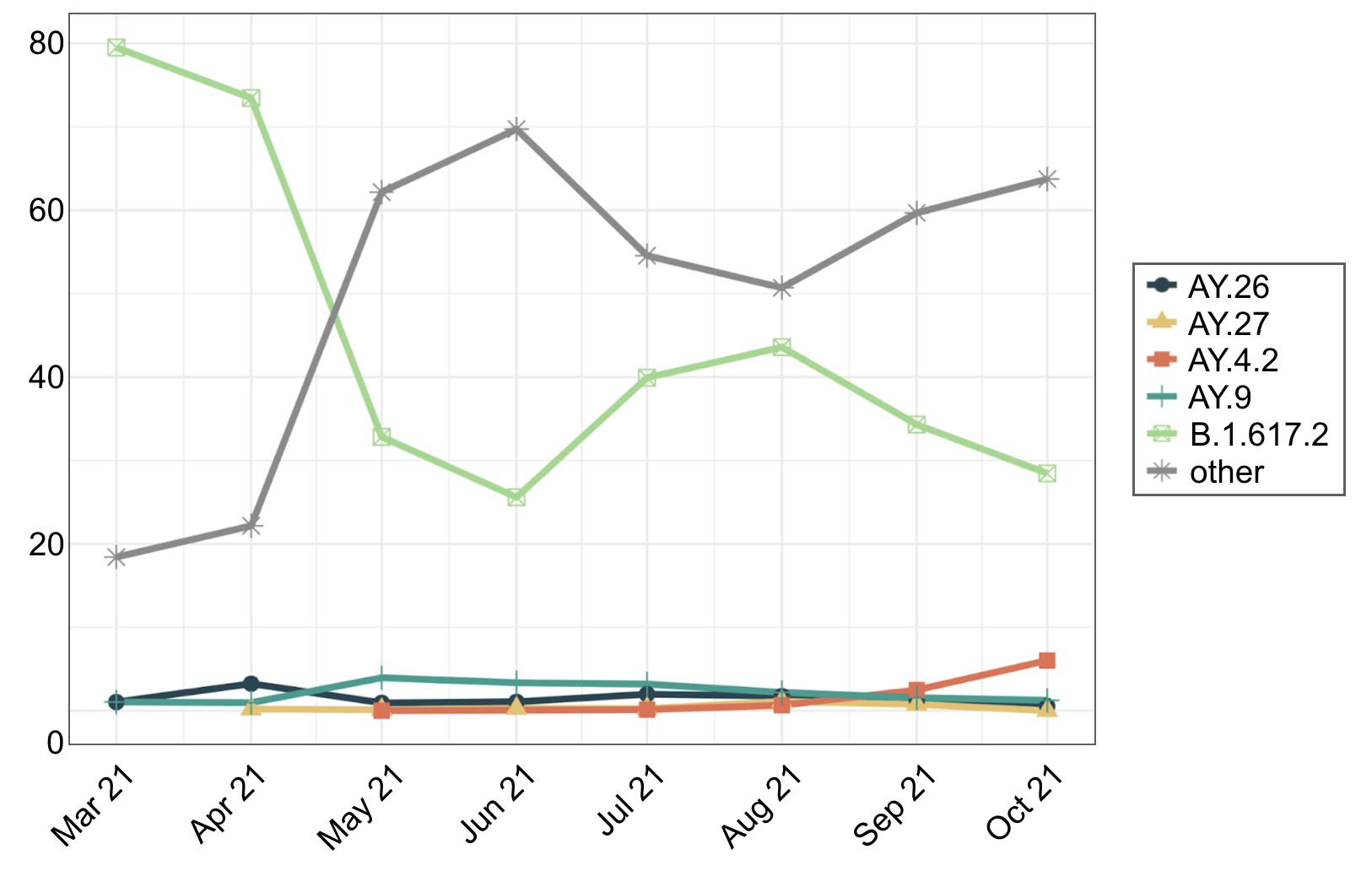

**Supplementary Figure 1. Temporal distribution of Delta sub-lineages.** Percentage of SARS-CoV-2 sequences designated as parental B.1.617.2 and its sub-lineages (designated as AY).

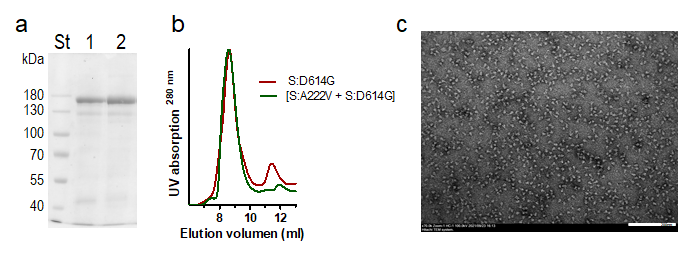

**Supplementary Figure 2. Protein purification and quality analysis. (a)** SDS-PAGE (Coomasie Staining) of the (1) S:D614G or (2) S:[A222V + S:D614G] spike variants. St, Pageruler Prestained Protein Ladder (Thermofisher Scientific) with masses (in kDa) indicated at the sides. (**b)** Size-exclusion chromatographic (SEC) profiles (UV absorption) of the indicated spike variants. (**c)** Electron micrograph of negatively stained particles from SEC fraction of D614G Spike protein, with scale bar = 200 nm.

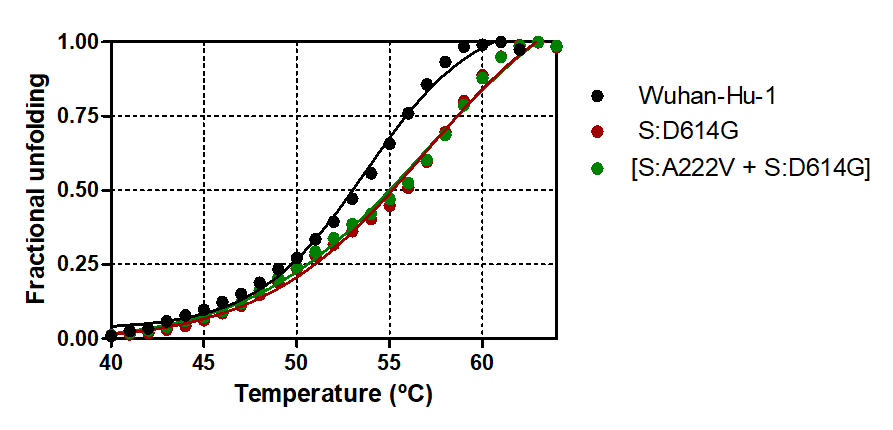

**Supplementary Figure 3.** **Thermofluor assays (means of three replicas) of the indicated SARS-CoV-2 S protein mutants.** Error bars are represented (too small to be seen).

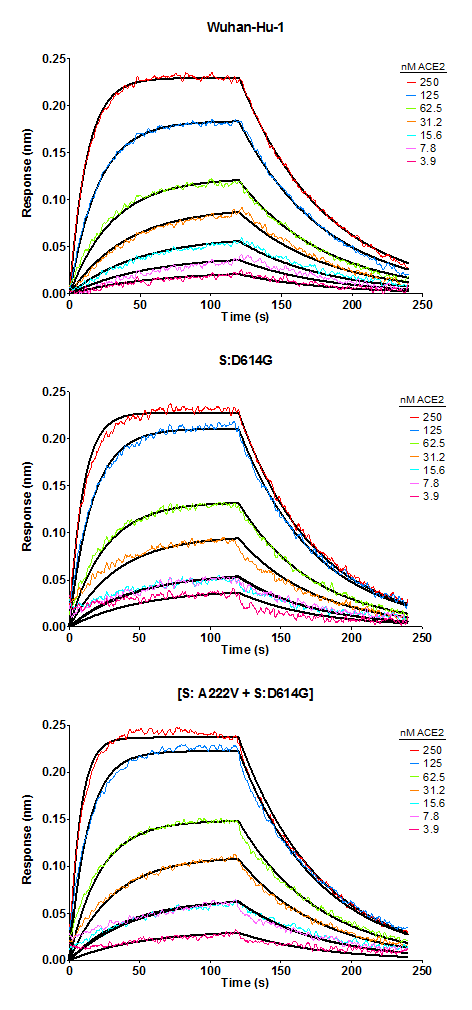

**Supplementary Figure 4. Kinetic analysis of spike interaction to ACE2 measured by biolayer interferometry (BLI).** Wuhan-Hu-1, S:D614G and [S:A222V + S:D614G]. The association of the different spike variants to ACE2 was carried out for 120s at various concentrations in a two-fold dilution series from 250 to 3.9 nM prior to dissociation for 120s. Curves were plotted using GraphPad Prism 6 for macOSX and fitting was performed using a 1:1 binding model in the data analysis HT software (Fortebio). Calculation of on-rates (K_on_), off-rates (K_off_) and affinity constants (K_D_) were computed using a global fit applied to all data. Raw data are coloured representations and fitting models are shown in black. Results are summarized in **Table 1** of the main text.

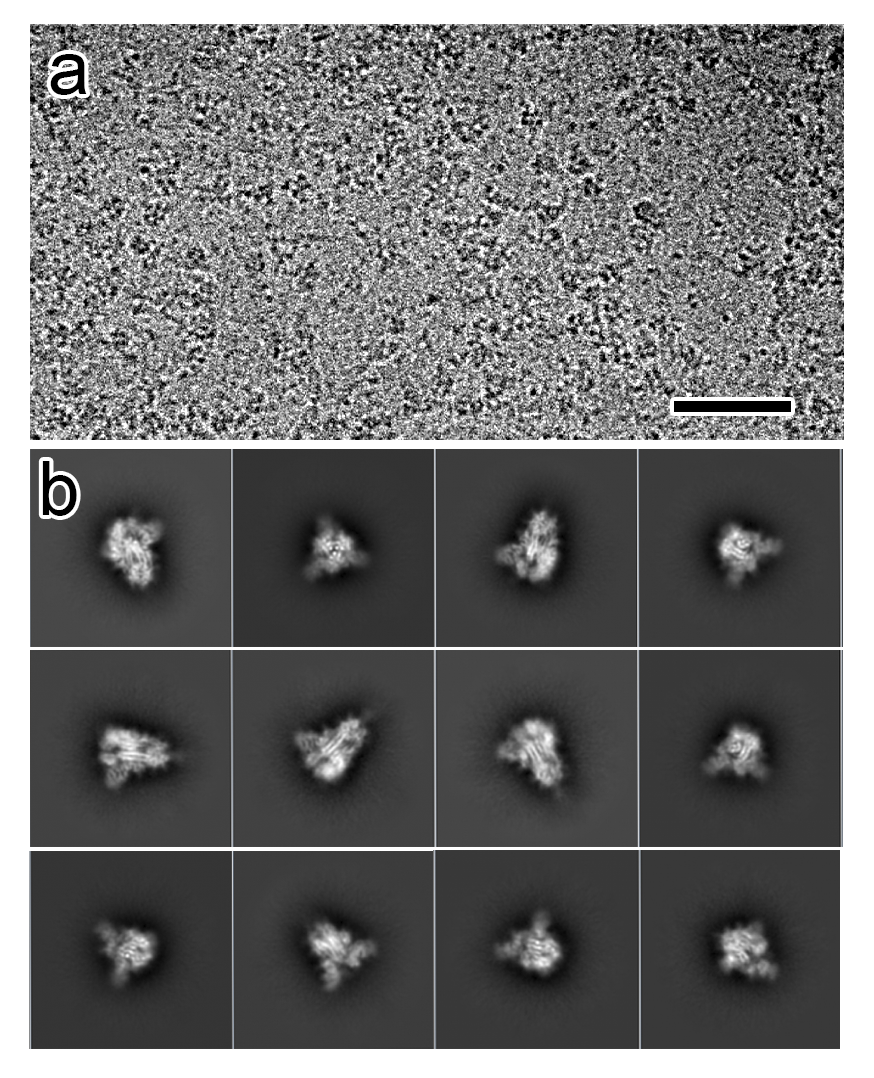

**Supplementary Figure 5. Cryo-EM data of [S:A222V + S:D614G]**. (**a**) Representative micrograph of the [S:A222V + S:D614G] mutant. Bar = 50 nm. (**b**) Set of representative side and top view class averages obtained after reference-free 2D classification of automatically picked and extracted particles.

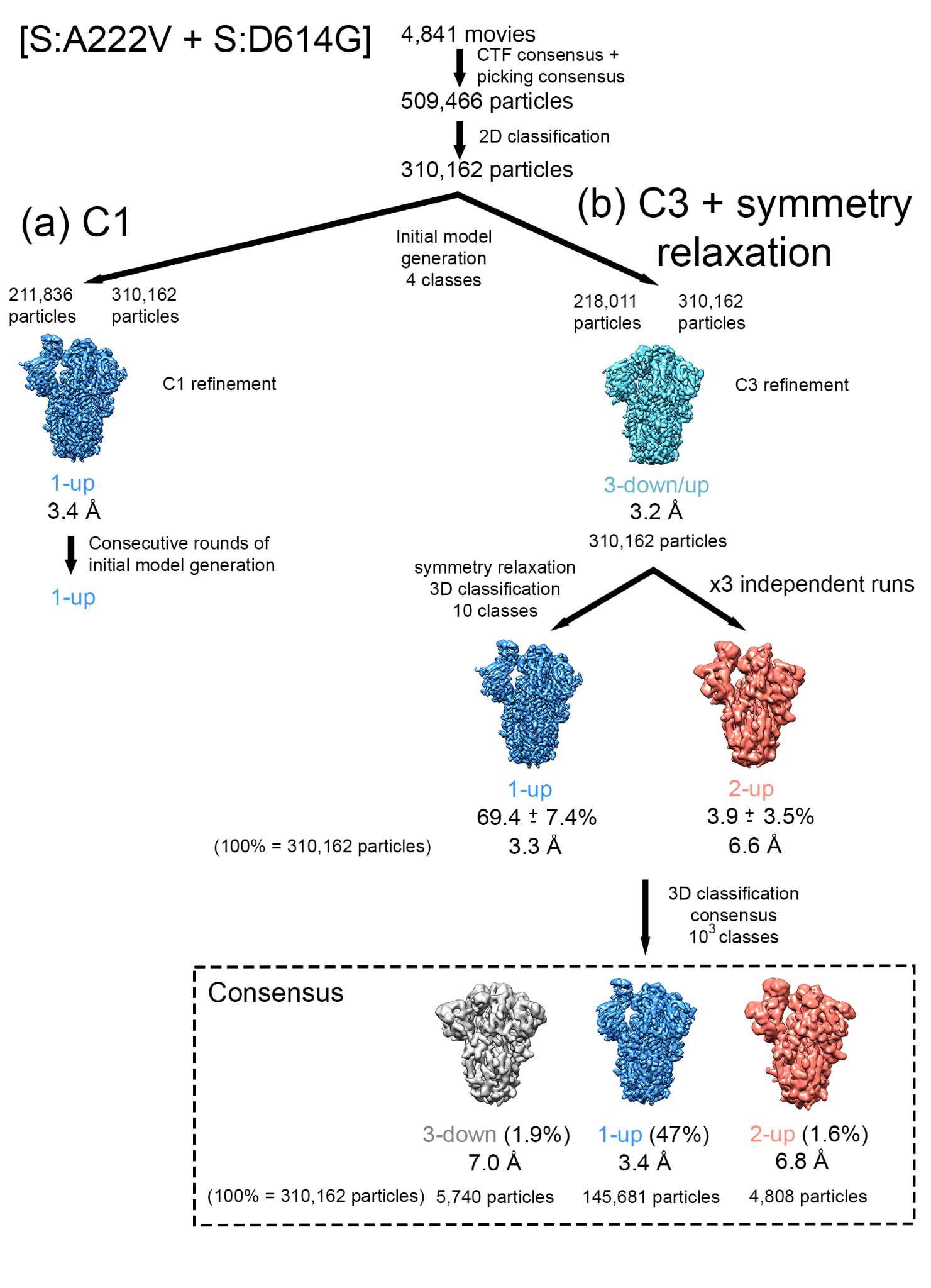

**Supplementary Figure 6. Cryo-EM image processing workflow for [S:A222V + S:D614G].** The initial dataset of 4,841 movies was subjected to motion-correction alignment, followed by CTF correction, automatic particle picking and subsequent extraction of 509,466 particles. After reference-free 2D classification, 310,162 particles were selected. Then, particles were subjected to initial model generation in 4 classes without (**a**) or with C3 symmetry imposition (**b**). In both image processing approaches, datasets considering unique or combined classes were refined to a resolution of 3.4 Å (**a**) and 3.2 Å (**b**). In (**a**), all of the maps obtained considering different classes and thus from different numbers of particles (211,836 or 310,162 particles) consisted of molecules with a 1-up conformation (blue). Consecutive rounds of initial model generation did not result in different RBD conformations, nor improvement of resolution. In (**b**), all of the maps obtained from different classes, each containing a different number of particles (218,011 or 310,162 particles), consisted of a 3-down/up mixed conformation (light blue) as a consequence of the symmetry imposition. The 310,162 particles dataset was then subjected to a first round of 3D classification with a symmetry relaxation implementation (**Methods**) in 10 classes and resulted in a main 1-up conformation (69%, blue) but also a minor 2-up conformation (7%, red). This 3D classification was repeated three times (mean and standard deviation are indicated) and subjected to the 3D classification consensus protocol. Note the power of the consensus approach (Sorzano et al. 2021), where a total of 10x10x10 = 1000 subclasses has been considered, selecting only those groups of particles that were always classified together (in other words, that they were stable clusters, coming together in three independent runs of classification). Naturally, results are considered to be stable only after consensus, presenting a % of occupancy much more reproducible than just after a first round. An initial model was generated for each selected class/conformation and subsequently refined to a resolution of 3.3-3.4 Å (1-up, blue), 6.6-6.8 Å (2-up, red) and 7 Å (3-down, grey). All of the maps shown were sharpened with DeepEMhancer (Sanchez-Garcia et al. 2021). Please note that the percentages shown do not add up to 100% (310,162 particles) in either analysis because some maps did not seem to be regular spikes and were assumed to contain junk particles.

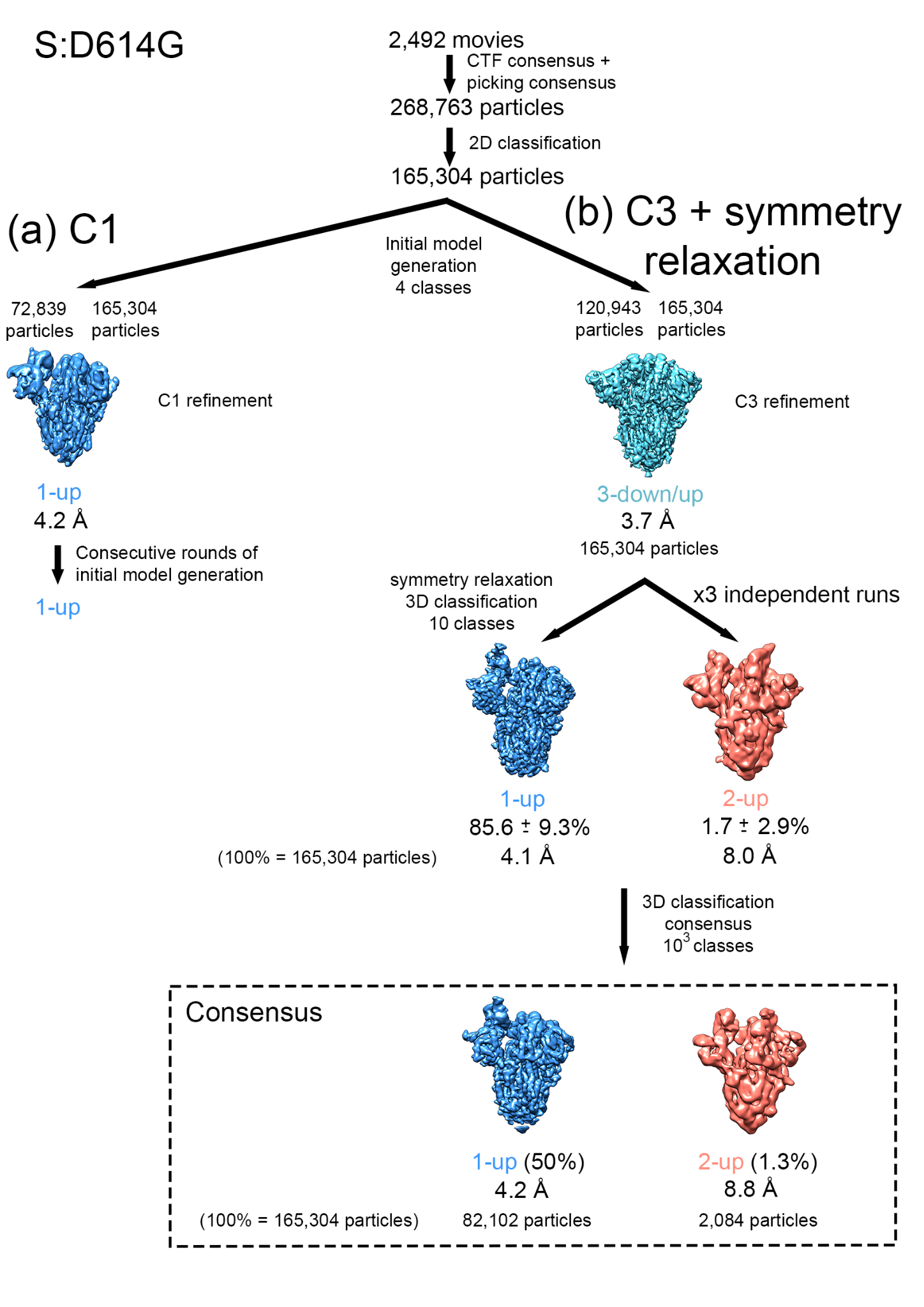

**Supplementary Figure 7.** **Cryo-EM image processing workflow for S:D614G.** The initial dataset of 2,492 movies was subjected to motion-correction alignment, followed by CTF correction, automatic particle picking and subsequent extraction of 268,763 particles. After reference-free 2D classification, 165,304 particles were selected. Then, particles were subjected to initial model generation in 4 classes without (**a**) or with C3 symmetry imposition (**b**). In both image processing approaches, datasets considering unique or combined classes were refined to a resolution of 4.2 Å (**a**) and 3.2 Å (**b**). In (**a**), all of the maps obtained considering different classes and thus from different numbers of particles (72,839 or 165,304 particles) consisted of molecules with a 1-up conformation (blue). Consecutive rounds of initial model generation did not result in different RBD conformations, nor improvement of resolution. In (**b**), all of the maps obtained from different classes and thus from different numbers of particles (120,943 or 165,304 particles), consisted of a 3-down/up mixed conformation (light blue) as a consequence of the symmetry imposition. The 165,304 particles dataset was then subjected to 3D classification with a symmetry relaxation implementation (**Methods**) in 10 classes and resulted in a main 1-up conformation (75%, blue) but also a minor 2-up conformation (5%). This 3D classification was repeated three times (mean and standard deviation are indicated) and subjected to the 3D classification consensus protocol as for the double mutant. An initial model was generated for each selected class/conformation and subsequently refined to a resolution of 4.1-4.2 Å (1-up, blue) and 8.0-8.8 Å (2-up, red). All of the maps shown were sharpened with DeepEMhancer (Sanchez-Garcia et al. 2021). Please note that the percentages shown do not add up to 100% (165,304 particles) in either analysis because some maps did not look like regular spikes and were assumed to contain junk particles.

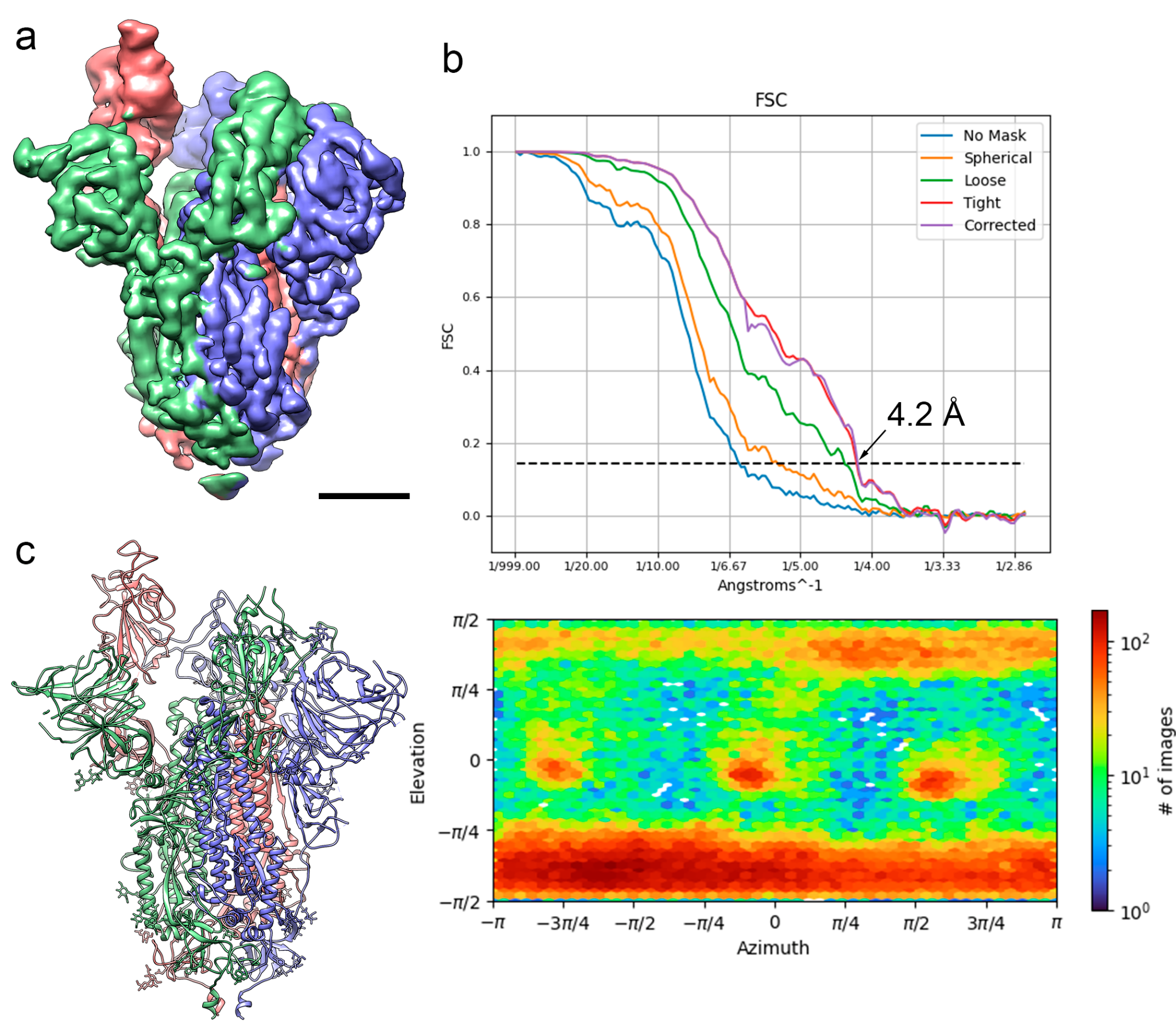

**Supplementary Figure 8. Cryo-EM of S:D614G. (a**) Side view of the cryo-EM density map of the S:D614G mutant obtained after consensus of three independent runs of symmetry relaxation, followed by initial model generation, refinement without symmetry imposition and deepEMhancer map sharpening, shown at 3 σ. Bar = 30 Å. (**b**) Fourier Shell Correlation (FSC) resolution curve (top) shown as the regular cryoSPARC global FSC resolution output, which includes no mask and different masks. Resolution based on the gold standard 0.143 criterion is 4.2 Å. Angular distribution coverage profile (bottom). (**c**) Atomic model of the D614G mutant shown as ribbon diagrams. S protein subunits are coloured in blue (A chain), red (B chain) and green (C chain). Glycan molecules are shown as stick diagrams and coloured according to their corresponding subunit.

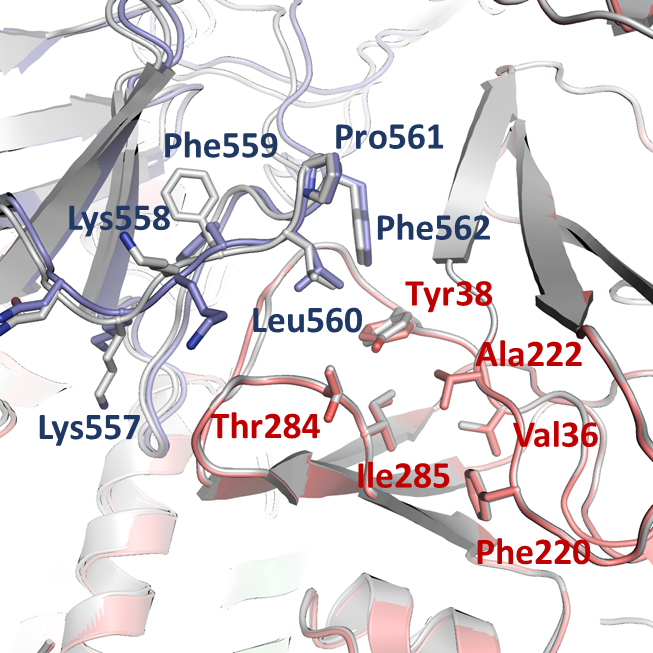

**Supplementary Figure 9. Comparison of the two structures at the region where S:A222V mutation is located.** The NTD_B_ and the CTD1_A_ of the double mutant [S:A222V + S:D614G] are coloured in salmon and blue respectively, while in the single mutant S:D614G, both subunits are coloured in grey. Side chains from residues surrounding the mutated residue are shown in sticks and indicated.

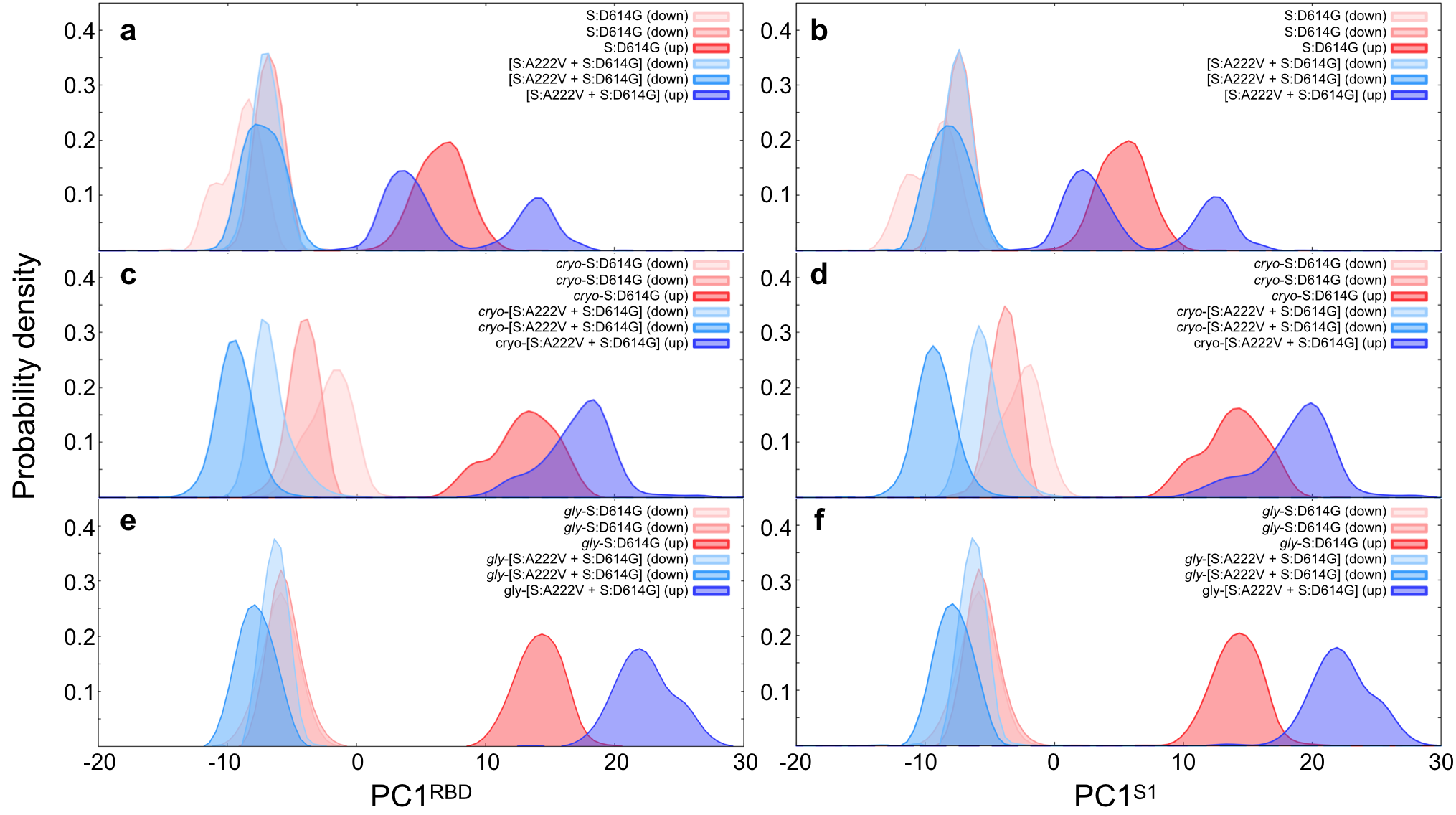

**Supplementary Figure 10. Continuous population densities along the PC1.** Continuous population densities along the first eigenvector for the PCA based on RBD (residues; 330-530; PC1^RBD^; **a, c, e**) and S1 (residues 30-650; PC1^S1^; **b, d, f**) of the MD-simulated 1-up (UDD) trimeric ensemble relative to the (**a**-**b**) glycan-free, (**c**-**d**) cryo-EM and (**e**-**f**) fully glycosylated S:D614G (as shades of red) and [S:A222V + S:D614G] (as shades of blue).

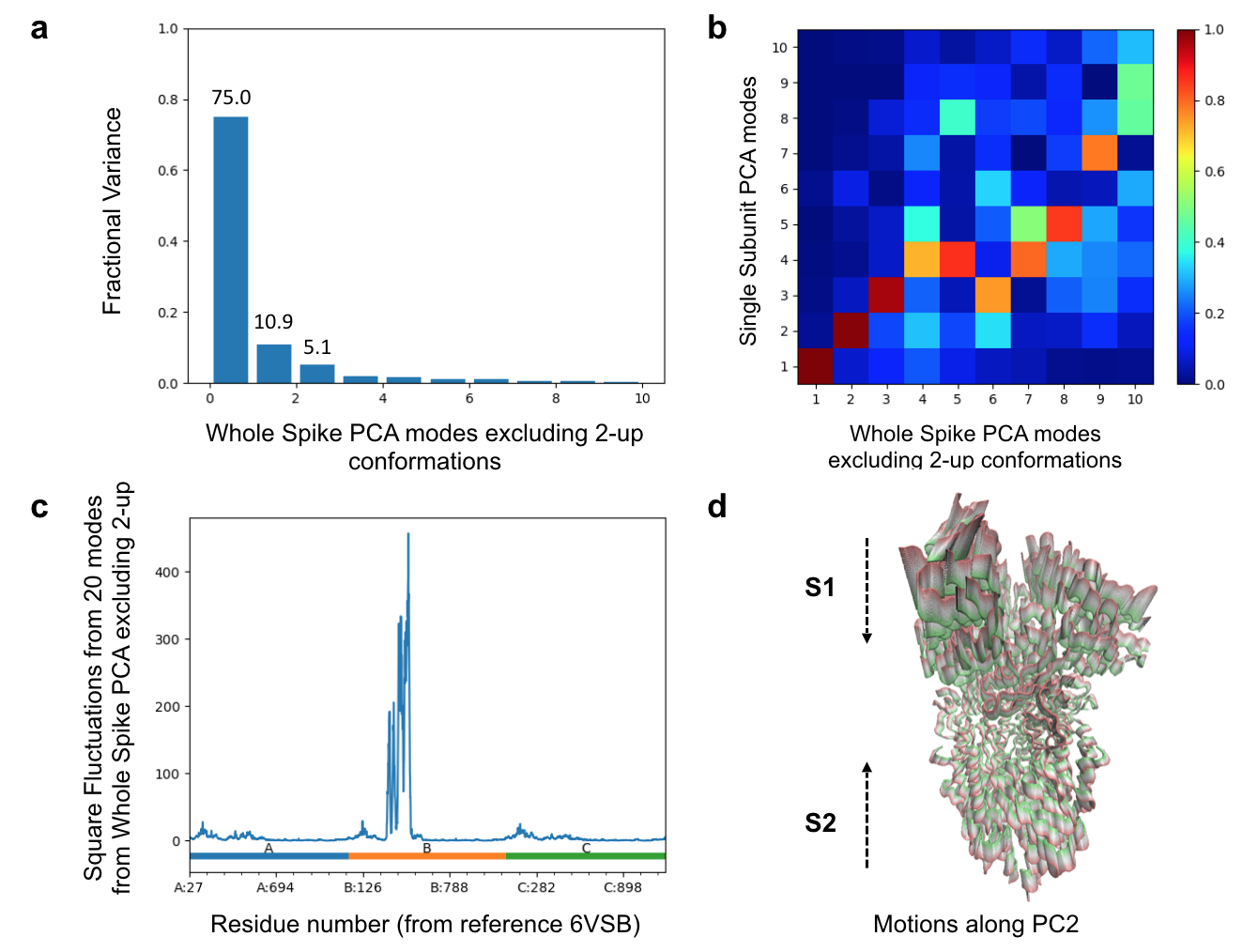

**Supplementary Figure 11. PCA of whole experimental S:D614G spike structures excluding 2-up conformations.** **(a)** Fractional variance contributed by each of the first 10 modes from the same whole spike PCA. Percentages (values multiplied by 100) are shown for the first three modes, which contribute more than 5 % (0.05) of the variation. **(b)** A matrix of correlation cosine overlaps comparing mode eigenvectors from a PCA performed using only the subunit whose RBD undergoes the transition to 1 up (ordinate) and that using the whole spike but excluding 2-up conformations (sliced to only include corresponding residues for the comparison; abscissa). The matrix is coloured from low overlaps in dark blue to high overlaps in dark red via cyan, green, yellow and orange as shown in the colour bar on the right. **(c)** Square fluctuations from the first 20 modes of variation from the same whole spike PCA weighted by the fractional variances. Residue numbers and chains are labelled as in our cryo-EM structures, allowing identification of motions of the dominant RBD (residues 330-530 of chain B) as well as other NTDs and RBDs. **(D)** Side view of the spike showing compression and stretching motions observed in whole spike PC2 using a range of states spanning 4 A from the average sone extreme in green (compressed state) to the other in red (stretched state).

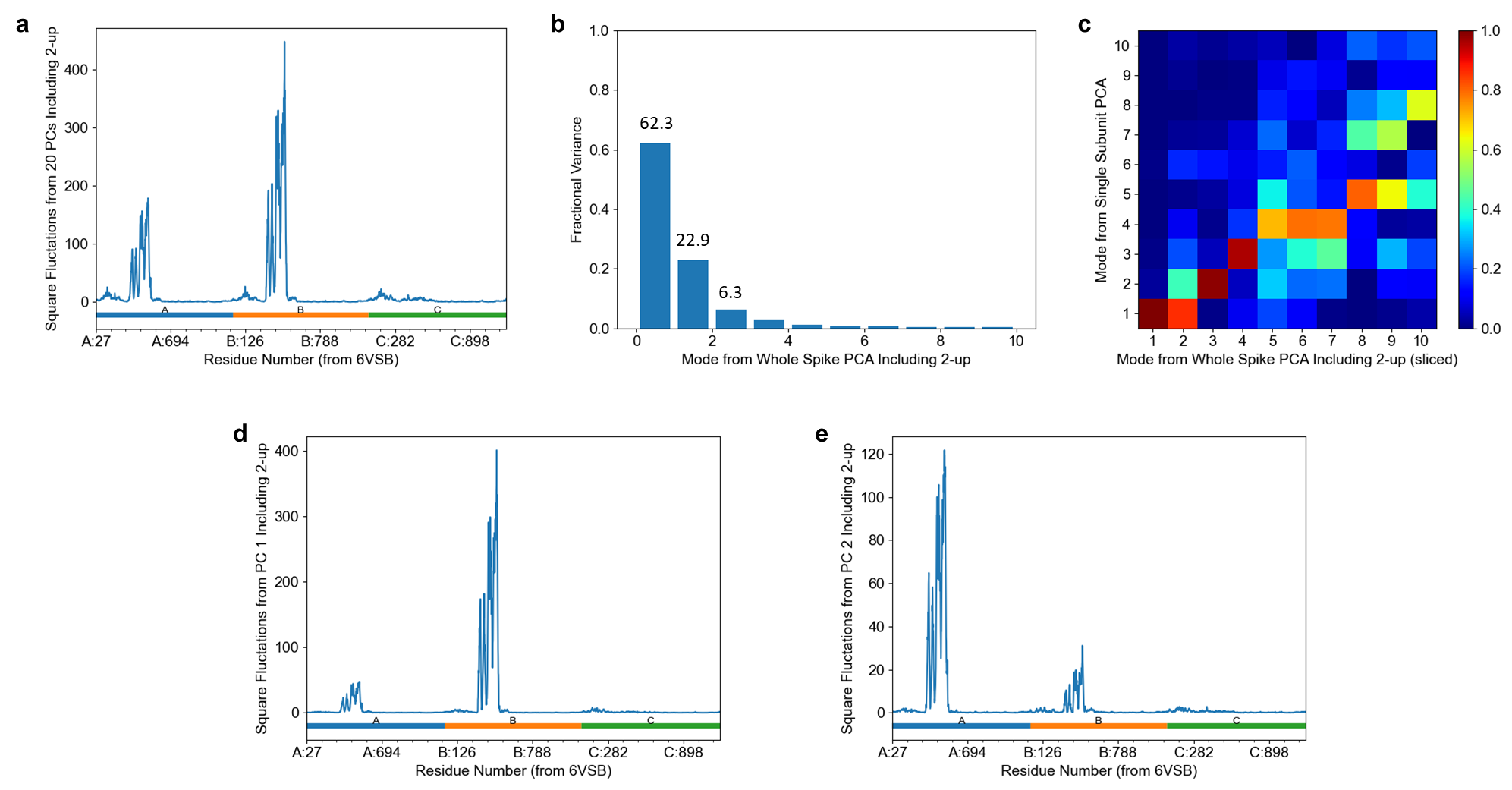

**Supplementary Figure 12.** **PCA results for whole S:D614G PDB structures including 2-up conformations.** **(a)** Square fluctuations from the first 20 modes of variation from the PCA from the whole spike excluding 2-up conformations weighted by the fractional variances. Residue numbers and chains are labelled as in our cryo-EM structures, allowing identification of motions of the dominant RBD (residues 330-530 of chain B) as well as other NTDs and RBDs. **(b)** Fractional variance contributed by each of the first 20 modes of the PCA from the whole spike excluding 2-up conformations. Percentages (values multiplied by 100) are shown for the first three modes, which contribute more than 5 % (0.05) of the variation. **(c)** A matrix of correlation cosine overlaps comparing mode eigenvectors from a PCA performed using only the subunit whose RBD undergoes the transition to 1 up (ordinate) and that using the whole spike including 2-up conformations in the covariance calculation (sliced to only include corresponding residues for the comparison; abscissa). The matrix is coloured from low overlaps in dark blue to high overlaps in dark red via cyan, green, yellow and orange as shown in the colour bar on the right. **(d-e)** Square fluctuations from PC1 **(d)** and PC2 **(e)**, showing dominance of the first RBD (chain B) and second RBD (chain A), respectively.

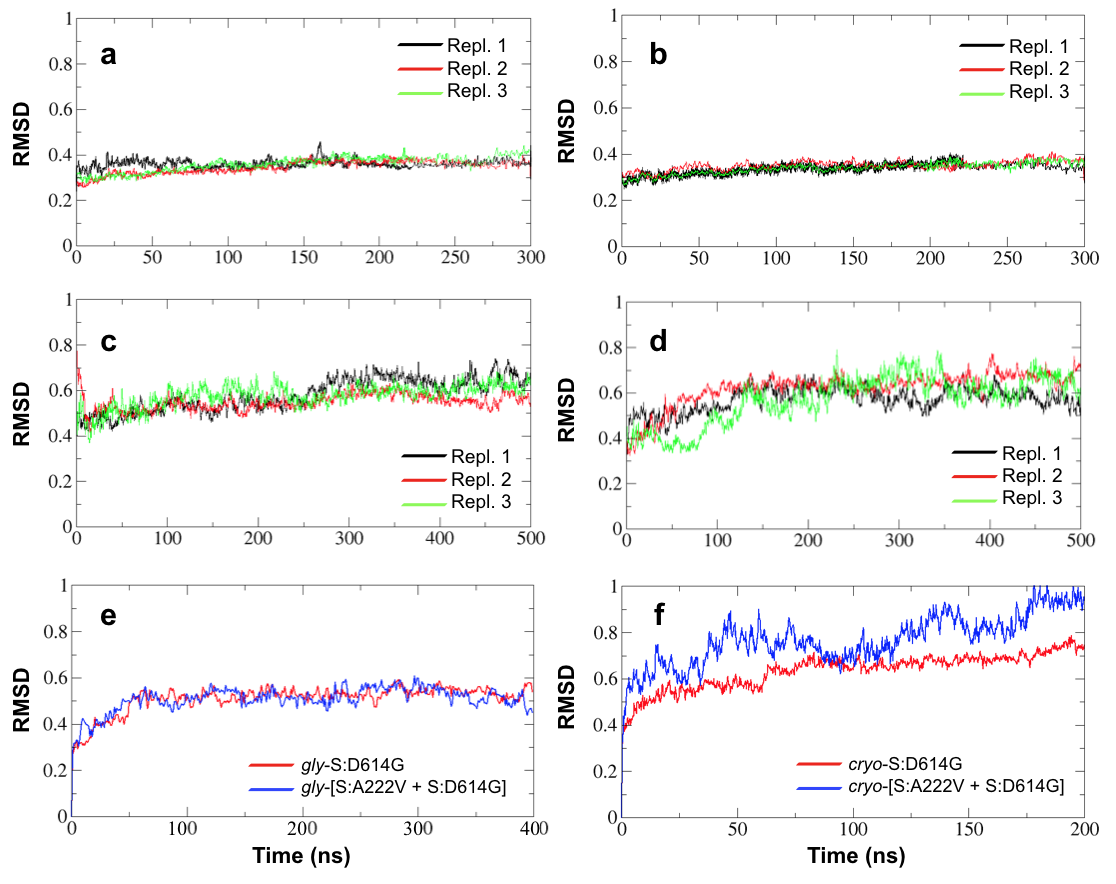

**Supplementary Figure 13. RMSD analysis.** Root-mean-square deviation (RMSD, nm) analysis of the MD trajectories (in triplicate) for the S:D614G (**a**, **c**) and [S:A222V + S:D614G] (**b**, **d**) mutants of the SARS-CoV-2 Spike mutants in the closed, 3-down (DDD; **a**, **b**) and open, 1-up (UDD; **c**, **d**) states respectively based on 6VXX and 6VSB. The same analysis for (**e**) the fully glycosylated mutants based on 6VSB and for (**f**) the (glycan-free) cryo-EM mutants in the open, 1-up state is also reported.

**Supplementary Video**

**Movie 1.** **Structural changes from S:D614G to [S:A222V + S:D614G], starting from cryo-EM maps, in two different orientations.** The subtle rearrangement of the RBDs and NTDs of subunits A and B of the spike is specifically highlighted. Position of residue 222 in each subunit is represented as a cyan sphere.

**Supplementary Tables**

**Supplementary Table 1.** Interaction between RBDs and NTDs domains from different subunits.

|  | **Interaction Surface (Å^2^)** | |
| --- | --- | --- |
|  | **[S:A222V + S:D614G]** | **S:D614G** |
| **Subunits A and B** | | |
| TOTAL | 1,011.3 | 882.4 |
| RBD_A_-NTD_B_ | 793.8 | 732.7 |
| RBD_A_-RBD_B_ | 217.5 | 149.7 |
| **Subunits B and C** | | |
| TOTAL (RBD_B_-NTD_C_) | 746.7 | 748.2 |
| **Subunits A and C** | | |
| TOTAL | 818.2 | 771.8 |
| NTD_A_-RBD_C_ | 744.2 | 717.7 |
| RBD_A_-RBD_C_ | 74 | 54.1 |

**Supplementary Table 2.** Detailed description of the 3-down (DDD) and 1-up (UDD) models of the SARS-CoV-2 spike mutants simulated in this study.

| **Glycan-free mutants from 6VXX/6VSB** | | | | |
| --- | --- | --- | --- | --- |
| **MODEL** | **N° atoms for the solvated systems** | **N° of TIP3P water molecules** | **N° of Na^+^ ions to neutralize** | **Simulated time**  **(ns)** |
| DDD_S:D614G_ |  |  |  |  |
| Replica 1 | 553,842 | 167,613 | 3 | 300 |
| Replica 2 | 553,842 | 167,613 | 3 | 300 |
| Replica 3 | 553,842 | 167,613 | 3 | 300 |
| DDD_[S:A222V + S:D614G]_ |  |  |  |  |
| Replica 1 | 553,860 | 167,613 | 3 | 300 |
| Replica 2 | 553,860 | 167,613 | 3 | 300 |
| Replica 3 | 553,860 | 167,613 | 3 | 300 |
| UDD_S:D614G_ |  |  |  |  |
| Replica 1 | 735,249 | 228,082 | 3 | 500 |
| Replica 2 | 735,249 | 228,082 | 3 | 500 |
| Replica 3 | 735,249 | 228,082 | 3 | 500 |
| UDD_[S:A222V + S:D614G]_ |  |  |  |  |
| Replica 1 | 735,258 | 228,079 | 3 | 500 |
| Replica 2 | 735,258 | 228,079 | 3 | 500 |
| Replica 3 | 735,258 | 228,079 | 3 | 500 |
| **Glycosylated mutants from 6VSB** | | | | |
| **SYSTEM** | **N° atoms for the solvated systems** | **N° of TIP3P water molecules** | **N° of K^+^/Cl^-^ ions to neutralize** | **Simulated time**  **(ns)** |
| *gly-*UDD_S:D614G_ |  |  |  |  |
| Replica 1 | 746,558 | 226,799 | 709/694 | 400 |
| *gly-*UDD_[S:A222V + S:D614G]_ |  |  |  |  |
| Replica 1 | 746,558 | 226,799 | 709/694 | 400 |
| **cryo-EM mutants of this work** | | | | |
| **SYSTEM** | **N° atoms for the solvated systems** | **N° of TIP3P water molecules** | **N° of K^+^/Cl^-^ ions to neutralize** | **Simulated time**  **(ns)** |
| *cryo-*UDD_S:D614G_ |  |  |  |  |
| Replica 1 | 826,453 | 257,517 | 767/767 | 200 |
| *cryo-*UDD_[S:A222V + S:D614G]_ |  |  |  |  |
| Replica 1 | 826,510 | 257,530 | 767/767 | 200 |

**Supplementary Table 3. Mutational free energy analysis.** Alchemical free energy calculations for the S:D614G and [S:A222V + S:D614G] mutants in the open, 1-up conformation based on 6VSB and our cryo-EM data. The obtained relative free energy values for the analysed systems indicate that the impact of the S:A222V mutation on the free energy of RBD opening is too small to establish a significant preference for either of the conformational states of the RBD.

| **Conformation** | ***gly*-UDD, S:A222V^a^** | ***cryo*-UDD, S:A222V^a^** | ***gly*-UDD, S:D614G^b^** |
| --- | --- | --- | --- |
| **Open** | 0.58 | 0.15 | 0.00 |
| **Closed (1)** | 0.45 | 0.00 | 4.15 |
| **Closed (2)** | 0.00 | 0.17 | 2.26 |

^a^ Relative values for S:A222V calculated on the background of S:D614G.

^b^ Relative values for S:D614G calculated on the background of Wuhan-Hu-1.

All values (in kcal/mol) are reported in relative terms (subtracting the lowest value). As a positive control, an analogous calculation using the wild-type (Wuhan-Hu-1) and a S:D614G mutant structure of glycosylated spike was performed to estimate the effect of the D614G mutation on the thermodynamics of RBD opening, and found a sizeable preference (of ca. 2-4 kcal/mol) for the open state.

**Supplementary Table 4.** Experimental structures of the SARS-CoV-2, S:D614G spike used in PCA analysis.

|  | **PDB ID** | **State^a^** | **CS Mutations** | **Proline Mutations** | **Chain Order** | **Reference** |
| --- | --- | --- | --- | --- | --- | --- |
| **1** | **6ZWV** | 3-down | - | - | BAC | Ke et al. 2020 |
| **2** | **7BNM** | 3-down | RSAS | PP | BCA | Benton et al. 2021 |
| **3** | **7BNN** | 1-up | RSAS | PP | BCA | Benton et al. 2021 |
| **4** | **7BNO** | 2-up | RSAS | PP | BCA | Benton et al. 2021 |
| **5** | **7KDI** | 3-down | - | - | BCA | Gobeil et al. 2021 |
| **6** | **7KDJ** | 1-up | - | - | BCA | Gobeil et al. 2021 |
| **7** | **7KE4** | 3-down | GSAS | - | BCA | Gobeil et al. 2021 |
| **8** | **7KE6** | 3-down | GSAS | - | BCA | Gobeil et al. 2021 |
| **9** | **7KE7** | 3-down | GSAS | - | BCA | Gobeil et al. 2021 |
| **10** | **7KE8** | 3-down | GSAS | - | BCA | Gobeil et al. 2021 |
| **11** | **7KE9** | 1-up | GSAS | - | BCA | Gobeil et al. 2021 |
| **12** | **7KEA** | 1-up | GSAS | - | BCA | Gobeil et al. 2021 |
| **13** | **7KEB** | 1-up | GSAS | - | BCA | Gobeil et al. 2021 |
| **14** | **7KEC** | 1-up, 1-I | GSAS | - | BCA | Gobeil et al. 2021 |
| **15** | **7KRQ** | 3-down | - | - | ABC | Zhang et al. 2021 |
| **16** | **7KRR** | 1-up | - | - | ABC | Zhang et al. 2021 |
| **17** | **7KRS** | 1-I | - | - | ABC | Zhang et al. 2021 |
| **18** | **7EAZ** | 1-up | GSAG | PP | ABC | Yang et al. 2021 |
| **19** | **7EB0** | 1-up | GSAG | PP | ABC | Yang et al. 2021 |
| **20** | **7EB3** | 1-up | GSAG | PP | ABC | Yang et al. 2021 |
| **21** | **7EB4** | 2-up | GSAG | PP | BCA | Yang et al. 2021 |
| **22** | **7EB5** | 2-up | GSAG | PP | ABC | Yang et al. 2021 |
| **23** | **7DX1** | 1-up | GSAS | PP | ABC | Yan et al. 2021 |
| **24** | **7DX2** | 1-up | -/digested | PP | ABC | Yan et al. 2021 |

^a^ “I” stands for Intermediate.

**Supplementary Table 5.** **Oligonucleotides used in this study.**

| Name | Sequence (5’-3’) |
| --- | --- |
| FW_ D614G_SPIKE | TCTTTACCAGGGCGTTAATTGTAC |
| RV_ D614G_SPIKE | ACAGCCACCTGGTTTGAC |
| FW_ A222V_SPIKE | GGGTTTTTCCGTACTAGAACCATTG |
| RV_ A222V_SPIKE | TGAGGTAAGTCGCGTAC |

**References**

Benton, D. J., A. G. Wrobel, C. Roustan, A. Borg, P. Xu, S. R. Martin, P. B. Rosenthal, J. J. Skehel, and S. J. Gamblin. 2021. “The effect of the D614G substitution on the structure of the spike glycoprotein of SARS-CoV-2.” *Proc Natl Acad Sci U S A.* 118 (9): e2022586118. DOI: 10.1073/pnas.2022586118.

Gobeil, S. M., K. Janowska, S. McDowell, K. Mansouri, R. Parks, K. Manne, V. Stalls, et al. 2021. “D614G Mutation Alters SARS-CoV-2 Spike Conformation and Enhances Protease Cleavage at the S1/S2 Junction.” *Cell Rep.* 34 (2): 108630. DOI: 10.1016/j.celrep.2020.108630.

Ke, Z., J. Otón, K. Qu, M. Cortese, V. Zila, L. McKeane, T. Nakane, et al. 2020. “Structures and distributions of SARS-CoV-2 spike proteins on intact virions.” *Nature.* 588 (7838): 498-502. DOI: 10.1038/s41586-020-2665-2.

Sanchez-Garcia, R., J. Gomez-Blanco, A. Cuervo, J. M. Carazo, C. O. S. Sorzano, and J. Vargas. 2021. “DeepEMhancer: a deep learning solution for cryo-EM volume post-processing.” *Commun Biol.* 4 (1): 874. DOI: 10.1038/s42003-021-02399-1.

Sorzano, C. O. S., A. Jiménez-Moreno, D. Maluenda, E. Ramírez-Aportela, M. Martínez, C. Cuervo, R. Melero, et al. 2021. “Image Processing in Cryo-Electron Microscopy of Single Particles: The Power of Combining Methods.” *Methods Mol Biol.* 2305:257-289. DOI: 10.1007/978-1-0716-1406-8_13.

Yan, R., Y. Zhang, Y. Li, F. Ye, Y. Guo, L. Xia, X. Zhong, X. Chi, and Q. Zhou. 2021. “Structural basis for the different states of the spike protein of SARS-CoV-2 in complex with ACE2.” *Cell Res.* 31 (6). DOI: 10.1038/s41422-021-00490-0.

Yang, T. J., P. Y. Yu, Y. C. Chang, and S. D. Hsu. 2021. “D614G mutation in the SARS-CoV-2 spike protein enhances viral fitness by desensitizing it to temperature-dependent denaturation.” *J Biol Chem.* 29 (4): 101238. DOI: 10.1016/j.jbc.2021.101238.

Yurkovetskiy, L., X. Wang, K. E. Pascal, C. Tomkins-Tinch, T. P. Nyalile, Y. Wang, A. Baum, et al. 2020. “Structural and Functional Analysis of the D614G SARS-CoV-2 Spike Protein Variant.” *Cell.* 183 (3): 739-751.e8. DOI: 10.1016/j.cell.2020.09.032.

Zhang, J., Y. Cai, T. Xiao, J. Lu, H. Peng, S. M. Sterling, R. M. Walsh, et al. 2021. “Structural impact on SARS-CoV-2 spike protein by D614G substitution.” *Science.* 372 (6541): 525-530. DOI: 10.1126/science.abf2303.
